# Portable Automated Rapid Testing (PART) for auditory assessment: Validation in a young adult normal-hearing population

**DOI:** 10.1101/2020.01.08.899088

**Authors:** E. Sebastian Lelo de Larrea-Mancera, Trevor Stavropoulos, Eric C. Hoover, David A. Eddins, Frederick J. Gallun, Aaron R. Seitz

## Abstract

This study aims to determine the degree to which Portable Automated Rapid Testing (PART), a freely-available program running on a tablet computer, is capable of reproducing standard laboratory results. Undergraduate students were assigned to one of three within-subject conditions that examined repeatability of performance on a battery of psychoacoustical tests of temporal fine structure processing, spectro-temporal amplitude modulation, and targets in competition. The Repeatability condition examined test/retest with the same system, the Headphones condition examined the effects of varying headphones (passive and active noise-attenuating), and the Noise condition examined repeatability in the presence of recorded cafeteria noise. In general, performance on the test battery showed high repeatability, even across manipulated conditions, and was similar to that reported in the literature. These data serve as validation that suprathreshold psychoacoustical tests can be made accessible to run on consumer-grade hardware and performed in less controlled settings. This dataset also provides a distribution of thresholds that can be used as a normative baseline against which auditory dysfunction can be identified in future work.

## I. Introduction

The assessment of auditory function in modern clinical audiology was translated from the laboratory in the middle of the previous century (Carhart & Jerger, 1959, Hughson & Westlake, 1944), and has remained focused on using pure-tone audiograms to evaluate audibility and speech tests to assess the ability to detect particular acoustical cues in speech (see CHABA, 1988). These clinical assessments are targeted at diagnosis of hearing impairment based on audibility and on an approach to rehabilitation that is largely defined by its reliance upon amplification via hearing aids. This focus on audibility and amplification has provided little incentive for clinical care to include the assessment and rehabilitation of supra-threshold auditory processing disabilities. As a result, there are very few tools, and even fewer protocols, available for the diagnosis and/or treatment of auditory difficulties that are not accompanied by losses of audibility. The diagnostic and rehabilitative approaches that do exist are regarded as specialized tools to be used by those clinicians who work with children or adults with suspected auditory processing disorders (APDs). There is a long history of clinicians and scientists using the term APD (e.g., Iliadou et al., 2018); yet, some clinicians and researchers are uncomfortable with the term due to the potential overlap of APD with language and cognitive dysfunction (e.g., Moore, 2018). The perspective taken by this study is that regardless of the clinical status of APD, it is undeniably the case that tests of auditory perceptual abilities (e.g., Moore et al., 2014; Eddins & Hall, 2010; Gallun et al., 2013) have the potential to shed light on complaints of hearing difficulties that are only weakly predicted by the audiogram or performance on clinical speech tests.

Clinically accessible tests of functional hearing are needed to better understand self-reported difficulties with auditory perception and poor performance on laboratory tests of auditory processing. These tests would need to be applied and validated across a population with diverse hearing abilities in order to clearly characterize which measures are most informative about the variety of hearing difficulties experienced by individual listeners or groups of listeners. Although a number of candidate tests have been developed and are relatively well studied in laboratory settings (e.g. Moore et al., 1987; Grose and Mamo, 2012; Bernstein et al., 2013; Gallun et al., 2014; Füllgrabe, Moore & Stone, 2015; Jakien et al., 2017; Hoover, Souza & Gallun, 2017; Hoover et al., 2019), very few of these tests have been translated into standard clinical practice. Those tests that have been translated into the clinic are generally only used by audiologists with expertise in APDs because the testing often requires specialized equipment or setup and a calibrated audiometer. Even when the tests are built into the audiometer, many audiologists have not received adequate training to feel comfortable administering, scoring, and interpreting the tests.

Tests that have moved successfully from the laboratory to the clinic include: the Staggered Spondaic Words test (SSW; Katz, 1962; Arnst, 1981), the Gaps in Noise test (GIN; Plomp, 1964; Green, 1971), the Masking Level Difference (MLD; Hirsh, 1948; Olsen, Noffsinger, Carhart, 1976), the Dichotic Digits Test (DDT; Broadbent, 1958; Musiek, 1983), the Listening in Spatialized Noise test (LISN; Cameron & Dillon, 2007; Glyde et al., 2013), the Frequency Patterns Test (FPT; Musiek & Pinheiro, 1987; Musiek, 1994), and the Dichotic Sentences Test (DST; Fifer et al., 1983). In addition, the Screening test for auditory processing (SCAN; Keith, 1995) is a battery of assessments that incorporates multiple auditory processing abilities. While these and other tests have been used successfully both in the laboratory and in the clinic to identify auditory processing dysfunction (e.g., Gallun et al., 2012; 2016; Hoover et al., 2017); however, none are portable, automated, or rapid—all require specialized equipment such as an audiometer, a trained audiologist to administer and score them, and most take 30 minutes to run and must then be scored by hand. The goal of this research project is to supplement these well-established tests with a low-cost, portable test system that could be used to administer a key set of basic auditory processing tests that is scored automatically and requires minimal clinical involvement. The assessments should each be rapid enough that clinicians and clinical researchers could tailor the length of the test battery to the time available. Moreover, portable automated rapid testing could play an essential role in gathering the datasets necessary to better characterize the auditory processing abilities and difficulties of individual listeners relative to the expected abilities of other listeners of a similar age with similar audiometric thresholds. Without this information, the clinician will continue to have difficulty appropriately identifying and remediating the auditory processing dysfunction they observe in their patients.

To address this gap, several state-of-the-art psychometric tests currently used in the laboratory to research central auditory processes have been translated into the application PART (Portable Automatic Rapid Testing) developed by the University of California Brain Game Center (https://braingamecenter.ucr.edu). PART can run both on mobile devices (e.g. iPad, iPhone, Android) and standard desktop computers (MacOS, Windows) and is currently freely available on the Apple App Store, the Google Play Store, and the Microsoft Store. PART has proven capable of accurately reproducing precise acoustic stimuli on an iPad (Apple, Inc., Cupertino, CA) with Sennheiser 280 Pro headphones (Sennheiser electronic GmbH & Co. KG, Wedemark, Germany) at output levels set by the built-in calibration routine (Gallun et al., 2018).

The psychophysical test battery evaluated here was designed to reflect a description of the central auditory system inspired by current research in psychoacoustics and auditory neuroscience (e.g., Stecker & Gallun, 2012; Bernstein et al., 2013; Depireux, Simon, Klein & Shamma, 2000). Although the specific tests described represent only a small subset of PARΓ’s functionality, the PART platform can facilitate a wider range of psychoacoustical tests. The full test battery assessed here comprises three sub-batteries, each with supporting evidence of clinical utility: temporal fine structure processing, spectro-temporal amplitude modulation, and targets in competition. These three groups of tests address different stages of auditory processing in the central nervous system that together mediate our ability to parse the auditory scene (Bronkhorst, 2015; Gallun and Best, in press).

Temporal Fine Structure (TFS) coding is assumed to rely upon the precision of phase-locking in populations of auditory nerve fibers responding to movements of the cochlear partition (Pfeiffer & Kim, 1975). The fine temporal information carried by the auditory nerve serves as the input to both the binaural system (see Stecker and Gallun, 2012) and the monaural pitch system (see Winter, 2005). Further refinement of this and other spectral and temporal information carried by the auditory nerve is responsible for the spectro-temporal modulation (STM) sensitivity observed in the inferior colliculus (Versnel, Zwiers & Opstal, 2009) and auditory cortex (Kowalsky, Depireux & Shamma, 1996). TFS sensitivity has been evaluated psychophysically using both monaural and binaural stimuli (Grose & Mamo, 2012; Gallun et al., 2014; Hoover et al., 2019). Neither the audiogram, nor most conventional speech tests, evaluate the detection of frequency modulation, or use spatialization of auditory signals; yet, it has been found that TFS measures are a good predictor of speech understanding in competition (Füllgrabe, Moore & Stone, 2015) and are suitable tests for age-related temporal processing variability (Grose & Mamo, 2012; Gallun et al., 2014; Füllgrabe, Moore & Stone, 2015). In this study, diotic frequency modulation was used to assess monaural TFS sensitivity, and dichotic frequency modulation was used to assess binaural TFS sensitivity. A temporal gap detection test (inter-tone burst delay) was also used to assess the sensitivity of temporal processes (Gallun et al., 2014). Because gap discrimination can be performed either using TFS information or by using envelope information carried by the auditory nerve (and refined by later processing), it is important to note that it is presently unclear which cue(s) are being evaluated, or even whether or not gap discrimination evaluates the same cues among different listeners. Nevertheless, these three tests have been proposed previously as measures of TFS with potential clinical utility (Hoover et al., 2019), and so, that category label is retained here for ease of reference.

Spectro-temporal modulation (STM) has been the increasing focus of laboratory studies as auditory cognitive neuroscience has revealed that cortical neurons are most sensitive to modulation of sound in both time and spectrum (Kowalsky, Depireux & Shamma, 1996; Theunissen, Sen & Doupe, 2000; Shamma, 2001; Schonwiesner & Zatorre, 2009). All natural sounds can be characterized as a pattern of spectro-temporal modulation (Theunissen, Sen & Doupe, 2000; Theunissen & Elie, 2014) and the relationship between sinusoidal spectro-temporal modulation and speech stimuli has been appreciated for some time (e.g., van Veen and Houtgast, 1985). This has led to a number of studies exploring sensitivity to spectral-, temporal-, and spectro-temporal modulation (STM) both for non-speech stimuli (e.g. Whitefield & Evans, 1965) and for speech stimuli (Bernstein et al., 2013; Mehraei et al., 2014; Venezia et al., 2019) as central processes that precede language understanding but require processing beyond basic audibility (Gallun & Souza, 2008). Studies using STM in participants with supra-threshold hearing loss have found that an extra 40% of the variance of speech-in-noise performance can be accounted for by these evaluations beyond the 40% accounted for by the audiogram alone (Bernstein et al., 2013; Mehraei et al., 2014). Thus, this study included tests for temporal-, spectral- and STM sensitivities, all of which are largely absent from the clinic.

Because the accurate identification of an acoustic target in competition is considered fundamental to auditory perception and scene analysis beyond peripheral audibility (Shinn-Cunningham, 2008; Moore, 2014; Bronkhorst, 2015), tests were included that assess the capacity of the system to select relevant information and suppress test-irrelevant interference. The notched-noise method (Patterson, 1976; Moore & Glasberg, 1990) evaluates the detection of a tone presented in competition with noise either with or without a spectral notch around the target frequency. This test allows the evaluation not only of peripheral frequency selectivity but also frequency processing efficiency (Patterson, 1976; Moore & Glasberg, 1990; Stone et al., 1992; Bergman et al., 1992). To address auditory scene analysis in the context of speech and binaural listening, spatial release from masking (SRM; Marrone et al., 2008; Gallun et al., 2013; Jakien et al., 2017; Jakien & Gallun, 2018) was assessed using the Coordinate Response Measure (CRM) corpus (Bolia et al., 2000). Following the methods of Gallun et al. (2013) speech understanding was assess both with speech maskers colocated with the target speech in simulated space, as well as with the maskers separated from the target by 45 degrees in simulated space. These tests independently assess speech understanding in competition under different stimulus conditions and the difference between the scores on the two provides a measure of the ability of an individual listener to benefit from spatial differences between target and masking stimuli.

The purpose of this study was to determine the degree to which this preliminary PART battery is capable of reproducing standard laboratory results in a population of young, normal-hearing adults. To this end, the reliability of threshold estimation (test-retest) and the degree to which estimates obtained from PART approximate those reported in the literature for the same tests were both evaluated. Additionally, to address the robustness of results to different listening conditions, we evaluated the extent to which test measures were consistent across the use of different headphone types and under different ambient noise conditions. Ultimately, the goal of this work is to generate a normative dataset that could be used in a range of contexts from research to the clinic.

To accomplish these goals, data were collected from young normal-hearing students from the University of California, Riverside campus under similar conditions to previous validation work from our group (Gallun et al., 2018) with repeated tests using circumaural headphones (Repeatability condition), with using both passive and active noise-attenuating headphones in a silent environment (Headphone condition), and in the presence of recorded cafeteria noise (Noise condition). First, the results addressing measurement reliability (test-retest) are presented. Second, the relation to the relevant literature is examined. Third, the effects of the experimental manipulations involving headphones and background noise are estimated. Overall, results show that PART produces repeatable threshold estimates consistent with those that have been reported previously in the laboratory across different listening conditions. These data serve as validation that accessible auditory hardware (consumer-grade tablet and headphones) can be used to test auditory function with sufficient precision to reproduce the thresholds obtained using laboratory-grade equipment. This dataset also provides a distribution of thresholds that can now be used as a normative baseline against which auditory dysfunction can be identified in future work.

## II. Methods

### A. Participants

Listeners were 150 undergraduate students from the University of California, Riverside (42 male, M age = 19.6 years, SD = 2.31 years), who received course credit for their participation. All participants reported normal hearing and vision, and no history of psychiatric or neurological disorders. They provided signed informed consent as approved by the University of California, Riverside Human Research Review Board. Consistent with our goal to evaluate “normal” auditory processing results presented in the main manuscript we rejected thresholds that deviated more than 3 SD from the mean on each assessment. A full data set with the thresholds of all participants is included in a form suitable for further analysis, and analyses and plots with the full dataset are included in the Supplemental Materials (see Fig S1 & table ST1).

### B. Materials

All procedures were conducted using standard iPad tablets (Apple, Inc., Cupertino, CA) running the PART Portable Automatic Rapid Testing (PART) application with stimuli delivered via either Sennheiser 280 Pro headphones (Sennheiser electronic GmbH & Co. KG, Wedemark, Germany), which are rated to have a 32 dB passive noise attenuation with an 8 Hz to 25 kHz frequency response, or Bose (active) noise cancelling Quiet Comfort 35 wireless headphones (Bose Corporation, Framingham, MA) set to the high noise cancelling setting. Output levels were calibrated for the Sennheiser headphones using an iBoundary microphone (MicW Audio, Beijing, China) connected to another iPad running the NIOSH Sound Level Meter App (SLM app; https://www.cdc.gov/niosh/topics/noise/app.html), as described in Gallun et al., 2018). The SLM app and iBoundary microphone system were calibrated with reference to measurements made with a Head and Torso Simulator with Artificial Ears (Brüel & Kjær Sound & Vibration Measurement A/S, Nærum, Denmark) in the anechoic chamber located at the VA RR&D National Center for Rehabilitative Auditory Research (NCRAR). Similar testing of the Bose system revealed that the method used, which did not involve changing the calibration settings when the headphones were changed, resulted in an overall reduction in the mechanical output level of 14 dB, but no distortions in the time or frequency domain. The levels described here and used throughout the study refer to the calibrated Sennheiser system.

### C. Procedure

In each session, participants sat in a comfortable chair in a double-walled sound-treated room and listened through a set of headphones connected to an iPad running PART. Tests were self-administered with text-based instructions delivered within the PART application. Responses were collected via digital-buttons presented on the iPad touch-screen. Most tasks employed a 2-cue 2-alternative forced choice (2-Cue 2-AFC) procedure where four intervals are presented in an audio-visual sequence with inter-stimulus-intervals (ISI) of 250ms (Fig. 1A top-left). The first and last stimuli were standard cues, and participants made a choice between the two alternatives presented in the second and third intervals (Fig. 1A top-right). Participants responded by touching the second or third square on the screen. The selected square then flashed either green (correct) or red (incorrect) as response feedback (Fig. 1A bottom) before proceeding to the next trial (1 sec ITI). This 2-cue 2-AFC task, which is identical to the one used in Souza et al. (2020), has the advantage that, unlike a two-interval or three-interval task, the target is always preceded and followed by a standard stimulus. This allows the task to be performed by comparing information either forward or backward in time. This is important as it is known that sensory comparisons are more difficult if they must be performed to a following standard rather than to a preceding standard, especially for older listeners (Gallun et al., 2012). A 2-cue 2-AFC design thus helps ensure that if in the future differences are found between the normative data reported here and data from other patient groups, that difference will be less likely to reflect the influences of attention or memory and more likely to reflect actual differences in the ability to make sensory comparisons. The one task that differed in procedure was the Spatial Release from Masking task (SRM) that uses a colored number grid to respond and has a fixed progression of difficulty (see details below).

**Fig 1.**
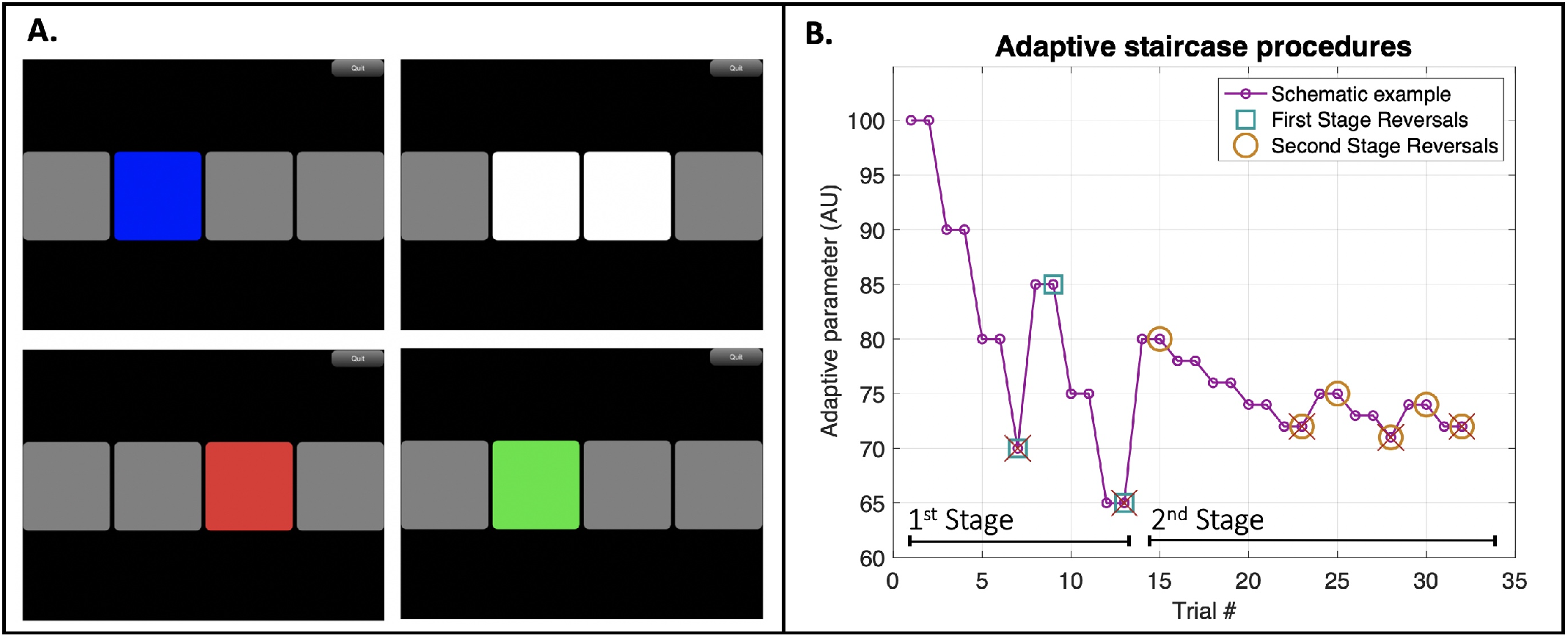
In A, each panel represents a screen-shot taken from PART while on a 2-cue, 2-alternative forced-choice test. Each box lit up sequentially in blue emitting a sound (top-left). After all intervals were played the 2 alternatives in the middle became available for response (top-right). Feedback is shown by color code (red = wrong; bottom panels). In B, we present a schematic example of the adaptive staircase procedures used. The difference in the magnitude of steps between staircase stages and the unequal step sizes going up/down can be easily observed in this example. Incorrect trials are marked with crosses and reversals are marked with either squares (1st stage) or circles (2nd stage). Arbitrary units were selected as adaptive parameter values for descriptive purposes only.

The tasks using the 2-cue 2-AFC procedure adjusted difficulty using a two-stage 2-down 1-up staircase procedure. The first stage used large steps for 3 reversals before moving on to the second stage that used smaller steps (1/5 the size of the first stage) and terminated after 6 reversals. Further, to help ensure that after incorrect responses participants were provided with easier exemplars, steps up were larger than steps down with a 2:1 step-size ratio in the Repeatability condition, and 1.5:1 step-size ratio in the Headphone and Noise conditions. Thresholds were estimated from the geometric mean of the second-stage reversals. A general schematic of the adaptive staircase procedures is included in Figure 1B. This combination of up-down rule and step-size ratio results in a threshold estimate that asymptotically targets the stimulus level corresponding to 81.7% correct for 2:1 and 77.5% correct for 1.5:1, comparable to a 79.4% targeted by a typical 3-down 1-up staircase with equal steps up and down (Levitt, 1971; García-Pérez, 2001; see section E of the results for the comparison across procedures). While unequal step sizes are common in audiometric testing (ANSI 3.21, 2004; ISO 8253-1, 2010), there are few who have followed the suggestion of García-Pérez (2011) in adopting the use of unequal step sizes when designing efficient staircase methods for clinical use. The goal is to minimize the influence of task and listener factors that can result in thresholds deviating from the asymptotic target point (García-Pérez, 1998; 2001). Designing optimal methods for the clinical translation of laboratory procedures is a continued area of research by our group (e.g., Hoover et al., 2019). The exploration of different ratios of unequal step sizes reported here represents an initial foray into this question.

Each session of the experiment began with a monitored screening test which presented 10 trials of a 2kHz tonal target signal at 45 dB SPL in the environmental settings relative to each condition. In cases where participants failed to respond accurately on at least 9 of the 10 trials, instructions were repeated in isolation to ensure that the task was properly understood. All of those participants who needed to restart the testing reported that they did not realize that the tone would be presented at a fairly low level. Once properly prepared for the stimuli to be at 45 dB SPL, all participants were able to detect the 2kHz tone with at least 90% accuracy. At this point, all participants moved on to complete two assessments involving the detection of the same 2kHz tone, but now presented in noise maskers with or without a spectral notch (described in detail below). Participants then were pseudo-randomly assigned to complete the remaining eight assessments (details described below) in three blocks of testing organized by test type: Temporal Fine Structure (TFS; 3 assessments); Spectro-temporal Modulation (STM; 3 assessments); and the second half of Targets in Competition (SRM; 2 assessments). All assessments were preceded by 5 non-adaptive practice trials at a high point in their respective staircase where target stimuli were easily detectable. Participants were encouraged to take small breaks between testing-blocks. All three test-blocks were given each session. The ten assessments in the testing block took around 5 minutes each, resulting in test sessions of around 50 minutes. The second session was always conducted on a different day, no longer than a week after the first. Test sessions involved up to three participants seated next to each other in a single room, listening and responding independently. In general, listeners received minimal instructions regarding the proper placement of the headphones and adherence to the brief written instructions automatically delivered by PART. The full verbal and written instructions given to each listener are provided in the Supplemental Materials.

### D. Stimuli

#### 1. Temporal Fine Structure (Fig. 2A)

a. *Temporal Gap* – This gap discrimination task (Gallun et al., 2014; Hoover et al., 2019) compares a target signal that consisted of a diotically presented temporal gap placed between two 0.5 kHz tone bursts of 4 ms played at 80dB SPL to standards that consisted of both tone bursts sequentially with no gap between them. The adaptive parameter was an inter-tone burst delay with an initial value of 20 ms. The staircase adapted on an exponential scale with first stage step-size (down) of 2^1/2^ and second stage step-size (down) of 2^1/10^ with a minimum of 0 ms and a maximum of 100 ms.
b. *Diotic Frequency Modulation* – This FM detection task (Grose & Mamo, 2012; Whiteford & Oxenham, 2015; Whiteford et al., 2017; Hoover et al., 2019) compares a target diotic frequency modulation of 2 Hz to standards that consisted of a pure tone carrier frequency randomized between 460 and 550 Hz, each presented at 75 dB SPL for 400 ms. Randomization of the carrier frequency of standards ensures that the test cannot be successfully conducted by a simple pitch cue. The adaptive parameter was modulation depth with an initial value of 6 Hz. The staircase adapted on an exponential scale with first stage step-size (down) of 2^1/2^ and second stage step-size (down) of 2^1/10^ with a minimum of 0 Hz and a maximum of 10 kHz.
c. *Dichotic Frequency Modulation* – This FM detection task (Grose & Mamo, 2012; Hoover et al., 2019) uses a stimulus first developed by Green et al. (1976), that create a continuously shifting interaural phase difference (IPD) in the target interval. The task compares a target signal consisting of a frequency modulation of 2 Hz that is inverted or is anti-phasic between the ears to standards that consisted of a pure tone carrier frequency randomized between 460 and 550 Hz, each presented at 75 dB SPL for 400 ms. The adaptive parameter was modulation depth (which determines the size of the IPD) with an initial value of 3 Hz. The staircase adapted on an exponential scale with first stage step-size (down) of 2^1/2^ and second stage step-size (down) of 2^1/10^ with a minimum of 0 Hz and a maximum of 10 kHz.

**Fig 2.**
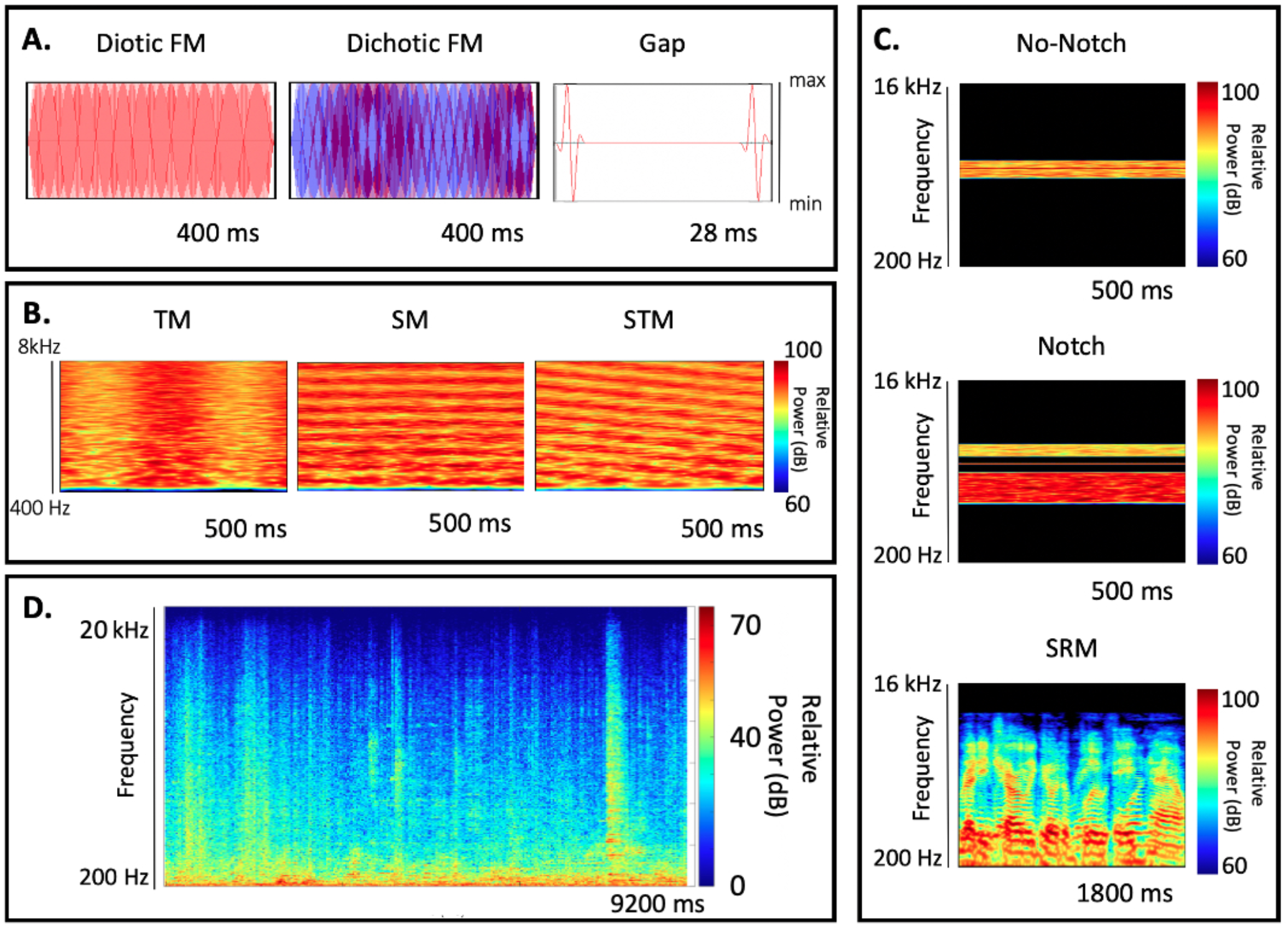
Visual descriptions of the stimuli employed for each assessment grouped by sub-battery are shown in sub-panels A-C. A representative nine-second segment of the cafeteria noise utilized for the Noise condition is shown in sub-panel D. The total recording had a duration of 33 minutes and was played in a continuous loop during testing.

#### 2. Spectro-Temporal Sensitivity (Fig. 2B)

All stimuli for these tasks involved a broadband noise that was either unmodulated (the standard) or modulated temporally, spectrally, or spectro-temporally, depending on the task (described below). The unmodulated standard consisted of flat-frequency broad-band noise with a frequency range of 0.4 to 8 kHz. Stimuli were generated in the frequency domain using the maximum number of components allowed by a 44.1 kHz sampling rate with random amplitude and phase values, presented at 65 dB SPL for 500 ms. Modulation was applied on a logarithmic amplitude scale (dB) and modulation depth was measured from the middle of the amplitude range to the peak amplitude. The stimuli were generated between trials using the algorithm developed by Stavropoulos et at. (under review).

a. *Temporal Modulation* – The TM detection task (Viemeister, 1979) compares a target with sinusoidal temporal amplitude modulation (AM) at a rate of 4 Hz to the unmodulated standard. The adaptive parameter was modulation depth in dB. The staircase adapted linearly in dB with first stage step-size (down) of 0.5 dB and second stage step-size (down) of 0.1 dB with a minimum of 0.2 dB Hz and a maximum of 40 dB.
b. *Spectral Modulation* – The SM detection task (Hoover, Eddins & Eddins, 2018) compares a target with a sinusoidal spectral modulation with random phase at a rate of 2 cycles per octave (c/o) to an unmodulated standard. The adaptive parameter was modulation depth in dB, which was adaptively varied as in the TM task.
c. *Spectro-Temporal Modulation* – This STM detection task (Bernstein et al., 2013; Mehraei et al., 2014) uses similar stimuli to the TM & SM tasks described above but compares a target with both 2 c/o spectral modulation and 4 Hz AM to standards that consisted of flat-frequency broadband noise. The resulting spectro-temporal modulation (STM) was randomly assigned to move upward or downward in frequency over time on each trial. The adaptive parameter was modulation depth in dB, which was varied as in the TM and SM tasks.

#### 3. *Targets in competition* (Fig. 2C)

a. *No-Notch Condition* – This abbreviated notch-noise method is adapted from Moore et al. (1987) and measures the ability of the listener to detect a target 2 kHz pure tone presented at 45 dB SPL in only one of the four intervals. The masking noise, which occurred on all intervals, consisted of 10,000 sinusoidal components distributed exponentially (−3 dB/octave) centered on the target frequency with a bandwidth of 800 Hz (1.6 to 2.4 kHz) presented for 500 ms. The adaptive parameter was the RMS level of the noise, measured in dB. The staircase started with a noise level of 35 dB SPL and adapted on a linear scale with first stage step-size (down) of 6 dB SPL and second stage step-size (down) of 2 dB SPL with a minimum of 25 and a maximum of 90 dB SPL.
b. *Notch Condition* – This condition was identical to the no-notch condition, with the exception that a spectral notch of 0.4 kHz was introduced, increasing the bandwidth of the masker such that it covered two frequency ranges: 1.2-1.6 kHz and 2.4-2.8 kHz, leaving a 0.4 kHz notch centered on 2 kHz, which was the frequency of the target to be detected. The adaptive parameter was the RMS masker level, which again had a starting value of 35 dB SPL. The staircase adapted in the same manner as in the no-notch condition. This condition is equivalent to a notch width of 0.2 times the center frequency of 2 kHz as described by Moore et al. (1987), and its difference in threshold with the no-notch condition can be taken as an index of frequency (spectral) resolution.
c. *SRM Colocated* – The three-talker speech-on-speech masking method of Marrone et al. (2008), as adapted for progressive tracking by Gallun et al. (2013), was used to measure the ability of listeners to identify keywords of a target sentence in the presence of two masking sentences. Using a color/number grid (4 colors by 8 numbers) participants identified two keywords (a color and a number) by selecting the position indicated by the keywords spoken by the target talker, which was a single male talker from the Coordinate Response Measure corpus (CRM, Bolia et al., 2000) presented from directly in front of the listener in a virtual spatial array. Target sentences all included the call-sign “Charlie” and two keywords: a number and a color. Targets were fixed at an RMS level of 65 dB SPL. The target was presented simultaneously with two maskers, which were male talkers uttering sentences with different call-signs colors and numbers in unison with each other and with the target. All three sentences were presented from directly in front of the listener (colocated). Progressive tracking included 20 trials in which the maskers progressed in level from 55 dB SPL to 73 dB SPL in steps of 2 dB every two trials as reported in Gallun et al. (2013), resulting in 2 responses at each of 10 target-to-masker ratios (TMRs). Threshold TMR was calculated following Gallun et al. (2013), by subtracting the number of correct responses from 10 dB, resulting in values between 10 dB for no correct responses to −10 dB for all correct responses. Negative TMR thresholds indicate threshold performance (roughly 50% correct) could be achieved when the target was at a lower level than the maskers, while positive thresholds indicate that the maskers needed to be lower in level than the target.
d. *SRM Separated* – The stimuli were identical to those in the colocated condition, with the exception that the maskers were presented from 45 degrees to the left and right of the target talker. Responses were again given in the context of a color/number grid (4 colors by 8 numbers) and participants had to select the position indicated by the target signal. Masker level again progressed every other trial from 55 dB SPL to 73 dB SPL in steps of 2 dB as reported in Gallun et al. (2013) and threshold TMR was again estimated by subtracting the number correct from 10 dB. The Spatial Release metric was estimated by subtracting the threshold in the Separated test from the threshold in the Colocated test, resulting in values between −20 dB and 20 dB, with 0 dB indicating no SRM, positive values indicating improvements in performance with spatial separation, and negative values indicating reduced performance with spatial separation.

### E. Experimental Design

The study consisted of 3-different conditions targeted to evaluate the repeatability of PART procedures in a variety of settings. These conditions were run sequentially on three different groups of participants.

1. ***Repeatability condition*** – The first 51 students enrolled were tested with Sennheiser 280 Pro headphones for both sessions and used 2:1 up/down step-size ratio in the staircase.
2. ***Headphone condition (in silence)*** – The next 51 participants enrolled were tested with different headphones (Sennheiser 280 Pro vs Bose Quiet Comfort 35) with the order counter-balanced between participants and used a 1.5:1 up/down step-size ratio.
3. ***Noise condition*** – The next 48 participants enrolled were tested using the same procedure as in the Headphone condition, but with recorded cafeteria noise played at 70 dB SPL. The noise was recorded in a local “coffee shop”, edited to remove silent gaps between recordings and transient recording noise at the beginning and ends of the recordings, and then bandpass filtered between 20 and 20,000 Hz. The coffee shop noise contained a large number of sound sources at all times, including both speech and environmental sounds. A spectrogram of a representative segment is shown in Fig. 2D. Sound files, after processing, were 33 minutes in duration and were played on a loop through two loudspeakers placed 30 cm apart from each other, and positioned in the center of the back of the test room, between 5 and 6 meters behind the three listeners.

## III. Results

Results are divided into sections for the purpose of clarity. First, the results for each test, session, and condition are presented (Section A). Then, issues of test-retest reliability are addressed (Section B) and the consistency of results for each test in comparison to previously reported measures are described (Section C). The effects of headphones and noise are addressed in Section D and the effects of staircase parameters in Section E. The full data set is provided in the Supplemental Materials for transparency and to encourage replotting, comparison with future and past data, and/or reanalysis.

### A. Overview

An overview of the results can be seen in Figure 3, which plots data for each test for each participant in each Condition and Session. This figure shows a high degree of repeatability between Session 1 and 2, and substantial overlap of performance between Conditions. This interpretation is consistent with summary statistics for each test shown as a function of Condition and Session in Table 1. Due to the high consistency between Conditions, the “main effects” were first analyzed by collapsing the data across Condition (Sections B, C) before addressing the effects of Condition (Section D). By combining data across conditions, a large normative dataset could be constructed, consisting of ~150 participants per test. Consistent with this goal of showing a normative sample, outlier rejection was performed by removing all data that exceeded three standard deviations from the mean for any condition of any task. The implications of this decision are addressed in the Discussion. Supplementary Materials are provided that demonstrate the robustness of results to different choices of outlier rejection as well as replotting data from Figure 3 with outliers included and labeled.

**Fig 3.**
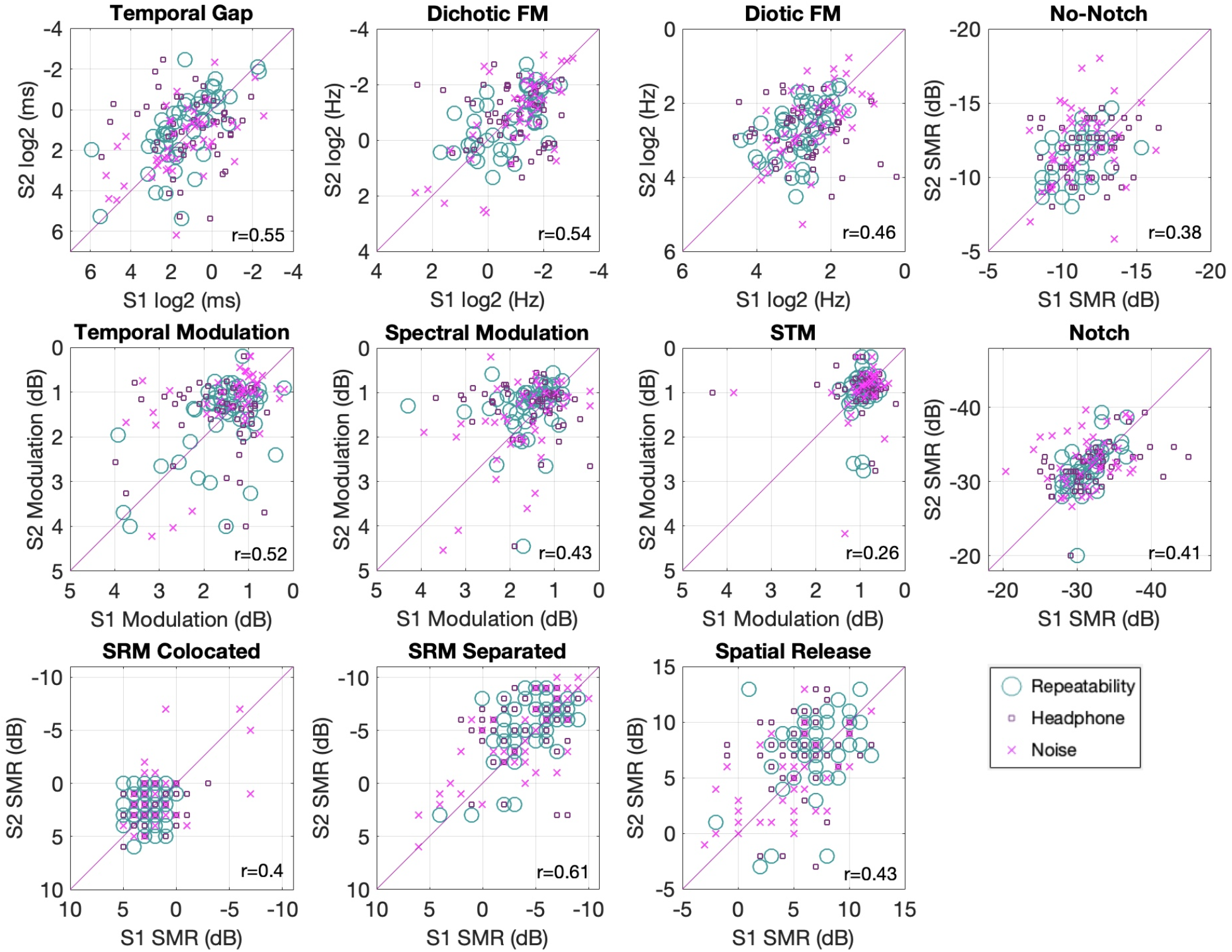
Scatter plots of Session 1 vs Session 2 for the 10 assessments. All axes are oriented to show better performance values away from the origin. Correlations are indicated in the lower right of each panel. Different markers illustrate the different conditions.

**Table 1.** Mean thresholds and standard deviations for the 10 assessments utilized plus the derived spatial release metric across all three conditions and their aggregate. Data are presented in PART’s native measurement units except for the targets-in-competition tests, which have been converted to TMR. The first row of each test shows session 1 and the second session 2 (S2).

| Test | *Repeatability*<br>*M (SD)* | *Headphone*<br>*M (SD)* | *Noise*<br>*M (SD)* | *All Cond.*<br>*M (SD)* | Units |
| --- | --- | --- | --- | --- | --- |
| Gap | 2.4 (2.9) | 2.17 (3.2) | 2.83 (3.41) | 2.46 (3.15) | Gap length |
| S2 | 1.8 (3.34) | 2.08 (3.04) | 3.008 (3.07) | 2.26 (3.18) | (ms) |
| Dichotic FM | 0.55 (2.09) | 0.54 (2.18) | 0.45 (2.44) | 0.51 (2.23) | Modulation Depth |
| S2 | 0.57 (2) | 0.47 (2.37) | 0.52 (2.77) | 0.52 (2.37) | (Hz) |
| Diotic FM | 7.07 (1.57) | 6.55 (1.88) | 5.48 (1.78) | 6.35 (1.75) | Modulation Depth |
| S2 | 6.71 (1.65) | 5.65 (1.76) | 5.82 (1.89) | 6.05 (1.77) | (Hz) |
| TM | 1.58 (.77) | 1.64 (.82) | 1.42 (.85) | 1.55 (.81) | Modulation depth |
| S2 | 1.58 (.85) | 1.46 (.85) | 1.21 (.84) | 1.42 (.85) | (dB) |
| SM | 1.57 (.61) | 1.55 (.75) | 1.71 (.82) | 1.61 (.72) | Modulation depth |
| S2 | 1.33 (.63) | 1.37 (.75) | 1.64 (.92) | 1.44 (.78) | (dB) |
| STM | 0.94 (.17) | .97 (.59) | .93 (.53) | 0.95 (.46) | Modulation depth |
| S2 | 0.97 (.49) | .98 (.55) | .94 (.62) | 0.96 (.55) | (dB) |
| No-Notch | -11.28 (1.38) | -11.82 (1.92) | -11.15 (2.14) | -11.43 (1.84) | Target-to-masker |
| S2 | -11.74 (1.71) | -12.88 (2.39) | -12.16 (2.48) | -12.26 (2.24) | ratio (dB) |
| Notch | -31.4 (2.27) | -32.26 (4.52) | -31.29 (3.82) | -31.67 (3.64) | Target-to-masker |
| S2 | -31.91 (3.02) | -32.71 (3.61) | -32.77 (3.31) | -32.44 (3.32) | ratio (dB) |
| SR Colocated | 2.33 (1.36) | 2.07 (1.67) | 1.92 (2.77) | 2.12 (1.96) | Target-to-masker |
| S2 | 2.01 (1.51) | 1.84 (1.96) | 1.34 (2.74) | 1.76 (2.08) | ratio (dB) |
| SR Separated | -4.58 (2.64) | -3.58 (2.93) | -3.57 (4.23) | -3.91 (3.32) | Target-to-masker |
| S2 | -5.36 (2.94) | -5.5 (2.64) | -4.19 (3.86) | -5.04 (3.2) | ratio (dB) |
| Spatial Release | 6.94 (2.78) | 5.66 (2.9) | 4.57 (3.72) | 5.8 (3.24) | Sep - Co |
| S2 | 7.34 (3.4) | 7.35 (2.71) | 4.67 (3.41) | 6.58 (3.37) | (dB) |

### B. Test-Retest Reliability

Test-retest reliability for the two sessions performed for each assessment in each experimental condition was evaluated using three metrics: Limits of Agreement (LoA, Altman & Bland, 1983; Bland & Altman, 1999), correlation, and t-tests. Each of these measures provides a different, but complementary, perspective on test reliability. The LoA analysis is considered a gold standard analysis as it provides information regarding both agreement and bias (e.g. systematic difference between sessions). Correlations are included to provide a measure of within subject consistency that ignores systematic effects of session, which can be important for research studies that seek to correct for effects of session. Lastly, t-tests were calculated as a function of session to help determine the reliability of effects of session.

#### 1. Test re-test reliability using limits of agreement (LoA)

For these data, LoA was considered to be a more informative measure of reliability than the normalized correlation, as between-subject variability for a sample that consisted solely of young listeners without hearing problems was anticipated to be small. Correlations, although more common in the auditory literature, are known to depend heavily on between-subject variability and measurement range (Altman & Bland, 1983; Bland & Altman, 1999). LoA plots for the estimated thresholds for each assessment are shown in Fig. 4 and the statistics are shown in Table 2. This analysis is based on the evaluation of performance across sessions (mean of test and re-test) as a function of their difference. LoA plots can be used to determine the extent to which learning effects are present—which would represent shifts towards better performance across sessions--, the region where 95% of the difference between test and re-test is expected to lie, and whether these statistics hold for different levels of performance (homoscedasticity).

**Fig 4.**
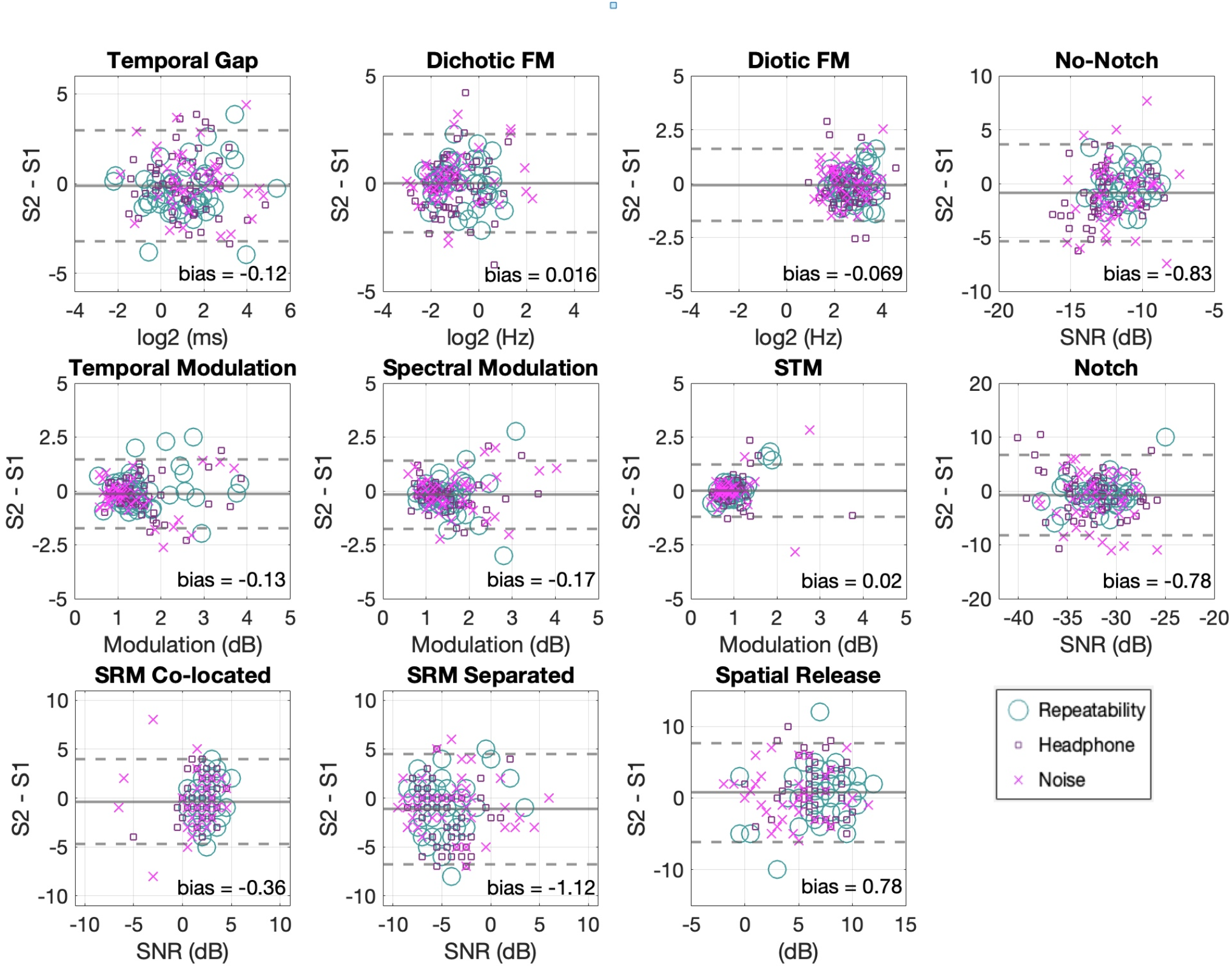
Limits of agreement of the estimated thresholds between sessions for all tests. The solid lines indicate the mean difference between sessions. Dotted lines indicate the 95% limits of agreement. Solid lines that fall below zero indicate better performance on session 2 (except the spatial release metric). Different markers illustrate the different conditions.

**Table 2.** Limits of agreement and within-subject significance testing for the 10 assessments utilized at two time points. Negative values on the bias column indicate better performance on the second session except on the Spatial Release metric, which is the only scale in which bigger magnitudes indicate better performance. * indicates significance at α = .05

| Test | Bias | Limits of Agreement | Units | $r(p)$ | $t(p)$ | Cohen's $d$ | df |
| --- | --- | --- | --- | --- | --- | --- | --- |
| Gap | -0.12 | [-3.2 to 2.96] | log2 (ms) | .55 (<.01)* | 0.94 (.34) | -0.07 | 145 |
| DichoticFM | 0.01 | [-2.2 to 2.2] | log2 (Hz) | .53 (<.01)* | -0.17 (.86) | 0.01 | 147 |
| DioticFM | -0.06 | [-1.73 to 1.59] | log2 (Hz) | .46 (<.01)* | 0.97 (.33) | -0.08 | 144 |
| TM | -0.13 | [-1.73 to 1.47] | M (dB) | .52(<.01)* | 1.94 (.054) | -0.15 | 145 |
| SM | -0.16 | [-1.75 to 1.41] | M (dB) | .58 (<.01)* | 2.46 (.01)* | -0.22 | 140 |
| STM | 0.01 | [-1.2 to 1.2] | M (dB) | .26 (<.01)* | -0.29 (.76) | 0.03 | 136 |
| No-Notch | -0.83 | [-5.3 to 3.6] | TMR (dB) | .38 (<.01)* | 4.37 (<.01) * | -0.4 | 144 |
| Notch | -0.77 | [-8.2 to 6.6] | TMR (dB) | .40 (<.01)* | 2.44 (.01)* | -0.22 | 142 |
| Colocated | -0.36 | [-4.7 to 3.9] | TMR (dB) | .39 (<.01)* | 1.95 (.052) | -0.17 | 142 |
| Separated | -1.12 | [-6.7 to 4.5] | TMR (dB) | .61 (<.01)* | 4.47 (<.01) * | -0.34 | 147 |
| SpatialR | 0.78 | [-6.1 to 7.6] | dB | .47 (<.01)* | -2.62 (<.01) * | 0.23 | 140 |

In order to facilitate visual inspection and comparisons across different tests, TFS tests were transformed from PART’s output units (either Hz or ms) to log2 units and target-in-competition tests were converted to target-to-masker ratios (subtracting the target level from PART’s masker level outputs). The mean across sessions is plotted on the x-axis to give a single point estimate for each participant in terms of their estimated threshold, thus showing between-subject variability of threshold estimation. The difference between sessions is plotted on the y-axis to give a single point estimate of the magnitude of deviation between sessions, thus showing within-subject variability of the estimated threshold. The mean of these differences is plotted as a straight line across the x-axis and its distance from zero (zero = perfect agreement) represents the main point estimate of the measurement’s systematic bias across sessions. The 95% limits of agreement (±1.96 *SD* (difference between sessions)) are plotted as dotted lines and indicate an estimate of the region in which we may expect to observe 95% of the within-subject, between-session differences of threshold estimation.

As can be observed in Fig. 4 and Table 2, the mean difference between sessions was close to zero in all of the tests, indicating little systematic bias. The 95% limits of agreement for the frequency modulation tests were at modulation rates of approximately ± 2 log_2_ (Hz) (or between 0.2 and 4.8 Hz). For the gap detection task, the LoA was approximately ± 3 log2 (ms) (or between 0.1 and 7.7 ms). The LoA for the modulation detection tasks were approximately ± 3 dB. For the speech tasks, the LoA are target-to-masker ratios of approximately ± 8 dB for the targets in competition tests.

Precise values are reported in Table 2. As can be seen in Fig. 4, the distribution of the threshold estimates had no salient asymmetries and session differences were similar across different levels of performance (symmetry along the abscissa). It is worth noting, however, that more spread can be identified at worse performance levels for some individuals in some tests. This applies both to the full set of data points in each plot and also to the subset of each showing the different conditions. There was little systematic bias between sessions (symmetry along the ordinate) suggesting similar measurement error for both sessions and that the poorer performance cases were expressed without a clear bias towards either session. This analysis demonstrates the range of alignment to be expected between different threshold estimates within subjects, and indicates that PART produces minimally biased estimates at the group level (see Table 2 for relevant statistics).

#### 2. Correlations Between Sessions

Table 2 shows statistics including the strength of association (Pearson r) between sessions. Significant correlations were observed for all the assessments. Overall, the relatively low correlation magnitudes reflect the warning of Altman and Bland (1983) that correlations are less informative to quantify reliability than LoA plots when performance is distributed across relatively narrow ranges of threshold estimates, as was to be expected for young listeners without hearing problems. This is particularly clear in the case of the STM assessment (in the same scale as the SM and TM assessments) where the range of threshold values obtained was quite restricted. In this context, the reduced between-subject variability in relation to a particular within-subject variance will have an impact on r-values decreasing their magnitude.

#### 3. Repeated-Measures T-tests

To supplement the LoA plots as a test for whether learning, or other factors, gave rise to systematic changes in performance, thresholds were compared between sessions across all three conditions using repeated-measures t-tests (see Table 2). While there were statistically significant differences between sessions in the spectral modulation detection test and the tone-in-noise tests, these changes were quite small, with magnitudes of less than 1 dB. The speech intelligibility test in the separated condition showed a significant difference of greater than 1 dB, which is consistent with the 1.58 dB difference previously reported by Jakien et al. (2017).

### C. Comparison with previously published results

While the above analyses demonstrate reliability of the test, it is possible that the rapid methods, the presence of noise, or the consumer-grade equipment would result in deviations from the results expected based on the published literature. This section thus compares the thresholds reported in Table 1 for all conditions averaged across sessions to those previously reported in the literature. This “grand mean” threshold estimate is included in Table 3. As outlier rejection was conducted by removing any points that fell more than three standard deviations above or below the mean, the number of outliers rejected and from which conditions is also reported, so that this can be considered in the comparisons. Overall, threshold estimates align with previous reports within 1.6 *SD* (see Table 3) and the number of outliers rejected was roughly consistent with the statistical expectation.

**Table 3.** Shows a summary of the similarities of the grand average thresholds estimated in the present study using PART and matched psychophysical tests from previous research. Plus or minus signs indicate values that are better or worse than previous reports respectively. The number of signs indicates increases in terms of SDs, one sign indicates < 1 SD and two signs indicate between 1 & 2 SD.

| *Assessment* | *Grand Average*<br>*M (SD)* | *Closest laboratory test* | *Distance in SD* |
| --- | --- | --- | --- |
| Gap | 2.36 (3.16) ms | Gallun et al., 2014 | – |
| Dichotic FM | 0.52 (2.3) Hz | Grose & Mamo, 2012;<br>Hoover et al., 2019 | – |
| Diotic FM | 6.2 (1.76) Hz | Grose & Mamo, 2012;<br>Hoover et al., 2019 | – – |
| TM | 1.49 (.83) M (dB) | Viemeister, 1979 | – |
| SM | 1.52 (.75) M (dB) | Hoover, Eddins & Eddins, 2005 | + + |
| STM | 0.95 (.51) M (dB) | Gallun et al., 2018 | - / + |
| No-Notch | -11.85 (2.09) TMR (dB) | Patterson, 1976 | – – |
| Notch | -32.06 (3.5) TMR (dB) | Patterson, 1976 | – – |
| SR Colocated | 1.94 (2.03) TMR (dB) | Jakien et al., 2017;<br>Gallun et al., 2018 | - / + |
| SR Separated | -4.47 (3.31) TMR (dB) | Jakien et al., 2017;<br>Gallun et al., 2018 | - / + |
| Spatial Release | 6.19 (3.32) (dB) | Jakien et al., 2017;<br>Gallun et al., 2018 | - / + |

#### 1. Temporal Fine Structure (TFS)

Sensitivity to temporal processing was assessed with three different tests; temporal gap detection, dichotic FM, and diotic FM. For temporal gap detection, 4 cases were rejected as outliers (Headphone condition 2; Noise condition 2), leaving threshold values that closely resemble those found in the literature (*M* = 2.36 ms, *SD =* 3.16). For example, Schneider et al. (1994) reported thresholds of 3.8 ms (right ear) and 3.5 ms (left ear) on average using 2kHz tone-bursts similar to the ones we used, however, their stimuli were delivered monaurally. Moreover, Hoover et al. (2019) reported thresholds of 1.45 ms using 0.75 kHz tone-bursts. Gallun et al. (2014) used the most similar stimuli (tone-bursts of 2 kHz) and obtained thresholds of 1.2 ms on average. All three of these studies used monaural presentation of their stimuli. Despite the differences in stimulus frequency and presentation style, all of these estimates lie within half a *SD* from the PART dataset. The fact that the published data report smaller thresholds and the second run appeared to produce smaller thresholds in this study suggest that the differences with the published literature might be removed by providing additional practice in the form of multiple measurements as opposed to the single track on each test session used here.

For the frequency modulated tests (dichotic & diotic FM), thresholds were higher than those previously reported in the literature. For the dichotic FM test, 2 cases were rejected as outliers (Repeatability condition 2). Thresholds in Hz (*M =* 0.52, *SD =* 2.29) are around 1 *SD* higher (on a logarithmic scale) than the 0.2 Hz found by Grose & Mamo (2012), the 0.15 Hz reported by Whiteford & Oxenham (2015), and the 0.19 Hz reported by Hoover et al. (2019).

For diotic FM, 5 cases were rejected as outliers (Repeatability condition 1; Headphone condition 3; Noise condition 1). Thresholds in Hz (*M =* 6.19, *SD =* 1.76), were about 2 *SD* higher than reports of Grose & Mamo (2012) of 1.9 Hz, Whiteford & Oxenham (2015) of .75 Hz, those of Moore & Sek (1996) of 1.12 Hz, and those of Hoover et al (2019) of 1.85 Hz, after conversion to Hz using the method of Witton et al. (2000) where appropriate. These differences in both FM tests are likely due to the difference in stimulus durations employed, which in the literature vary between 1000 ms (Moore & Sek, 1996) and 2000ms (Whiteford & Oxenham, 2015). Here, the duration was set to 400 ms. This choice was based on the results of Palandrani et al. (2019), who showed that FM detection thresholds improve with the square-root of stimulus duration. This predicts diotic FM thresholds of 3.6 Hz for the listeners here, if durations that were comparable to Grose & Mamo (2012) had been used. Even after this correction, however, the thresholds obtained were about 1 *SD* worse on average (on a logarithmic scale) after conversion to Hz using the method of Witton et al. (2000) where appropriate. As with the temporal gap, it would not be surprising if repeated testing resulted in reduced thresholds, more similar to those reported in the literature.

#### 2. Spectro-Temporal Modulation (STM)

Sensitivity to spectro-temporal modulation was assessed with three different tests; spectro-temporal modulation (STM), spectral modulation (SM) and temporal modulation (TM). It is difficult to make exact comparisons with previously reported results in the literature without making a variety of transformations and ignoring several differences in methodology. The most important issue is the measurement of modulation depth. Measurement depends on the scale (log or linear), the reference points (peak-to-valley or peak-to-midpoint), and the order in which the modulation operations are performed among other factors (see Isarangura et al., 2019). In this case, PART generated stimuli that were modulated on a logarithmic amplitude scale (dB), with modulation depth measured from the middle of the amplitude range to the peak amplitude. This differs from the method used by others, such as Hoover et al. (2018), who measured applied modulation that was sinusoidal on a dB scale but measured the amplitude as the difference from the maximum to the minimum, rather than the midpoint. Still more different was Bernstein et al. (2013), who applied sinusoidal modulation on a linear scale and also measured the modulation depth from the maximum to the minimum. When the amplitude scale is linear, threshold is expressed by transforming the modulation depth (m), which varies between 0 and 1, into dB units using the value 20 times log(m), which means that a fully modulated signal has a value of 0 dB, and a modulation depth of 0.01 has a value of −40. These differ from the values used to express threshold using a log amplitude scale, and thus Equation 1 (below) was used to convert the thresholds obtained with PART to 20log(m) dB units as detailed in Isarangura et al. (2019). Equation 1 accounts for both the use of a linear amplitude scale and for measurement of the amplitude from maximum to minimum. This provides a single metric that can be used to compare the temporal, spectral and spectro-temporal modulation thresholds obtained here to a subset of the data reported in the literature. It should be noted that there are other differences in signal generation, presentation methods, adaptive tracking, and training of participants that also may have influenced thresholds.

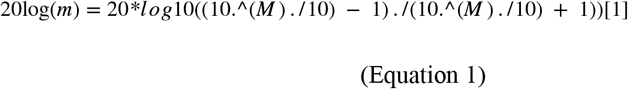

For STM at 4 Hz and 2 c/o, 13 cases were rejected as outliers (Repeatability condition 2; Headphone condition 3; Noise condition 8). STM thresholds obtained (*M =* 0.95 (M) dB, *SD =* 0.51) were converted using equation 1 to (*M =* −20.003 20log(m) dB, *SD =* 3.63) and closely resembled those previously reported in the literature. They were within a *SD* from those reported by Gallun et al. (2018) for five different testing sites (range −21.74 to −18.42 dB) and for Chi et al. (1999) (−22 dB). The obtained thresholds for STM are about 2 *SD* better than those reported by Bernstein et al. (2013) (−14 dB). It is unclear why outlier rejection was higher in this task than in others, but this may indicate that this is an ability on which some listeners are particularly poor. It is worth noting that the Supplemental Materials, where these statistics are reported without outlier rejection, still shows better performance than in the literature.

For SM at 2 c/o, 9 cases were rejected as outliers (Repeatability condition 1; Headphone condition 4; Noise condition 4). SM modulation depth thresholds (*M =* 1.52 dB, *SD =* 0.75) were converted using equation 1 to (*M =* −16.19 20log(m) dB, *SD =* 4.21) were better by about 1 *SD* than those reported by Hoover Eddins & Eddins (2018) (−11.08 dB), and those reported by Davies-Venn, Nelson & Souza (2015) (about −11 dB). These differences might be due to differences in modulation depth generation patterns or modulation depth metrics employed (see Isarangura et al., 2019). Further, stimulus parameters like those of the noise carrier bandwidth or presentation level, and test parameters such as tracking procedure varied across studies and so might account for the slight differences found. One reason to suspect that these methodological differences influenced performance is the fact that our listeners often outperformed the more practiced listeners in the other studies. Again, a higher number of outliers were observed, but as can be seen in the Supplemental Material, this did not account for the better performance in this study.

For TM at 4 Hz, 4 cases were rejected as outliers (Repeatability condition 1; Noise condition 3). TM thresholds (*M =* 1.49 (M) dB, *SD =* 0.83) are converted using equation 1 to (*M =* −16.63 20log(m) dB, *SD =* 4.61) were within half a *SD* of those reported by Viemeister (1979) of −18.5 dB for four observers.

#### 3. Target Identification in Competition

##### Tone Detection in Noise with and without a Spectral Notch

These tests evaluated the ability to detect a 2-kHz pure tone in competition with broad-band noise either overlaying the target signal (no-notch condition) or with a 400-Hz spectral notch or protective region without noise (notch condition). We rejected 5 cases as outliers (Headphone condition 1; Noise condition 4) from the No-Notch test and obtained thresholds of *M =* −11.85, *SD =* 2. In the case of the Notch test, we rejected 7 cases (Headphone condition 1; Noise condition 6) and obtained thresholds of *M =* −32.06, *SD* = 3.5. The notched-noise procedure has been widely used for the analysis of frequency selectivity in the cochlea (see Moore, 2012). Because of this, the emphasis of the literature has been on calculating detailed information about the shape of the auditory filter, and specific thresholds associated to each condition are typically not reported. However, Patterson (1976) reported an average distance between the equivalent of our no-notch and notch conditions of about 24 dB for four participants, which is comparable to the mean distance we obtained here of 20.20 dB (*SD* = 2.9) where some of our participants performed in the same range. This similarity and the high test-retest reliability of this test suggests that learning plays a small role in the ability to perform this test and that reliable estimates can be obtained with very few trials.

##### Speech-on-speech Competition

These tests evaluated the discrimination of speech in the face of speech competition using variants of the Spatial Release from Masking (SRM) test described by Gallun et al. (2013). Two conditions were used, one where the speech-based competition was colocated in virtual space with the target speech (colocated) and one where the speech-based competition was located ±45 degrees away from the target (separated) in simulated space. All values are reported in target-to-masker ratio (TMR) dB units. In the case of the colocated condition, 7 cases were rejected as outliers from the Noise condition. Interestingly, these cases are mainly due to performance that was better than average for more than 3 *SD* (see Supplemental Materials Fig S1), which has been observed previously for the occasional younger listener with normal hearing. Colocated thresholds (*M* = 1.94 dB, *SD* = 2.03) closely resemble those reported by Gallun et al. (2018) across two testing sites (1.85 & 1.96 dB). Performance was slightly worse than predicted by the normative functions of Jakien & Gallun (2018), which are based on linear regression to the data from a variety of listeners varying in age and hearing loss. For a 20 year old with a pure-tone average (PTA) of 5 dB HL, which seems appropriate for this sample, colocated thresholds averaged across two runs are predicted to be 1.2 dB, which is within 1 *SD* of what is observed.

In the Separated condition, 2 cases were rejected as outliers (Repeatability condition 1; Noise condition 1). Separated thresholds (*M* = −4.47 dB, *SD* = 3.31) closely resembled those reported by Gallun et al. (2018) across two testing sites (−4.33 & −4.62 dB), all of which were higher on average than the predictions of the equation of Jakien & Gallun (2018), which predicts a threshold of −6.7 dB, which is still within 1 *SD* of those observed.

The difference between the separated and the colocated conditions, a metric indicating the spatial release from masking effects, showed spatial release values (*M* = 6.19 dB, *SD* = 3.32) that again closely resembled the ones reported by Gallun et al. (2018) across two testing sites (6.19 & 6.57 dB) and were within 1 *SD* of the predictions of the regression equation of Jakien & Gallun (2018), which predicts 8.3 dB. Of note, the spatial release from masking magnitudes reported here are smaller than those reported for other similar tests already used in the clinic like the LiSN-S (Cameron & Dillon, 2007) which do not use synchronized concurrent masking and thus allow for better ear glimpsing (Bringart & Iyer, 2012) to contribute to the effect of release from masking (Glyde et al., 2013).

### D. The Effects of Headphones and Noise

To address the effects of headphone types with and without noise-attenuation technology and external noise conditions, “main effects” were evaluated by collapsing across tasks. To do so, composite scores were computed by normalizing each individual assessment relative to its mean and standard deviation (a z-score transform), calculating a z-score for each listener in each assessment, and then averaging these normalized values across the 10 assessments for each participant. Z-scores and composite scores are included in the data set that is available as part of the Supplemental Materials.

LoA plots, Pearson correlations, and t-tests were then computed for the composite scores. Results are reported for each condition separately, divided by headphone type. To test differences across experimental manipulations, a mixed-model Analysis of Variance (ANOVA) was used to compare composite scores across the factors of interest. The data for each condition as a function of test type are available in the Supplemental Materials.

The internal reliability of the composite score was assessed by calculating Cronbach’s α, which gave a score of 0.75. This indicates that the composite has strong internal reliability and thus it can be appropriate to use it as a summary score. Of note, the composite score is not an attempt to reduce central auditory processing to a single construct. Rather, this measure is intended to address the effects of these experimental manipulations in an efficient manner across all the assessments in the battery.

Fig. 5 shows the 95% limits of agreement for the standardized composite scores of the whole sample across three experimental conditions (panel on the left). This analysis showed close to zero bias (< 0.01), and limits of agreement of [−0.87, 0.88] which indicate that 95% of repeated estimates of the composite score of the battery used in this study are expected to lie within 1 SD from each other in young listeners without hearing problems. In addition, Fig. 5 also shows a scatterplot of session 1 vs 2 for the composite scores (panel on the right). This composite showed stronger association between sessions than each of the individual assessments (*r* = 0.65 *p* < .001, [95% *CI =* 0.55, 0.73]) and represents an alternative estimate to the LoA regarding the reliability of the battery as a whole and not of its individual assessments.

**Fig 5.**
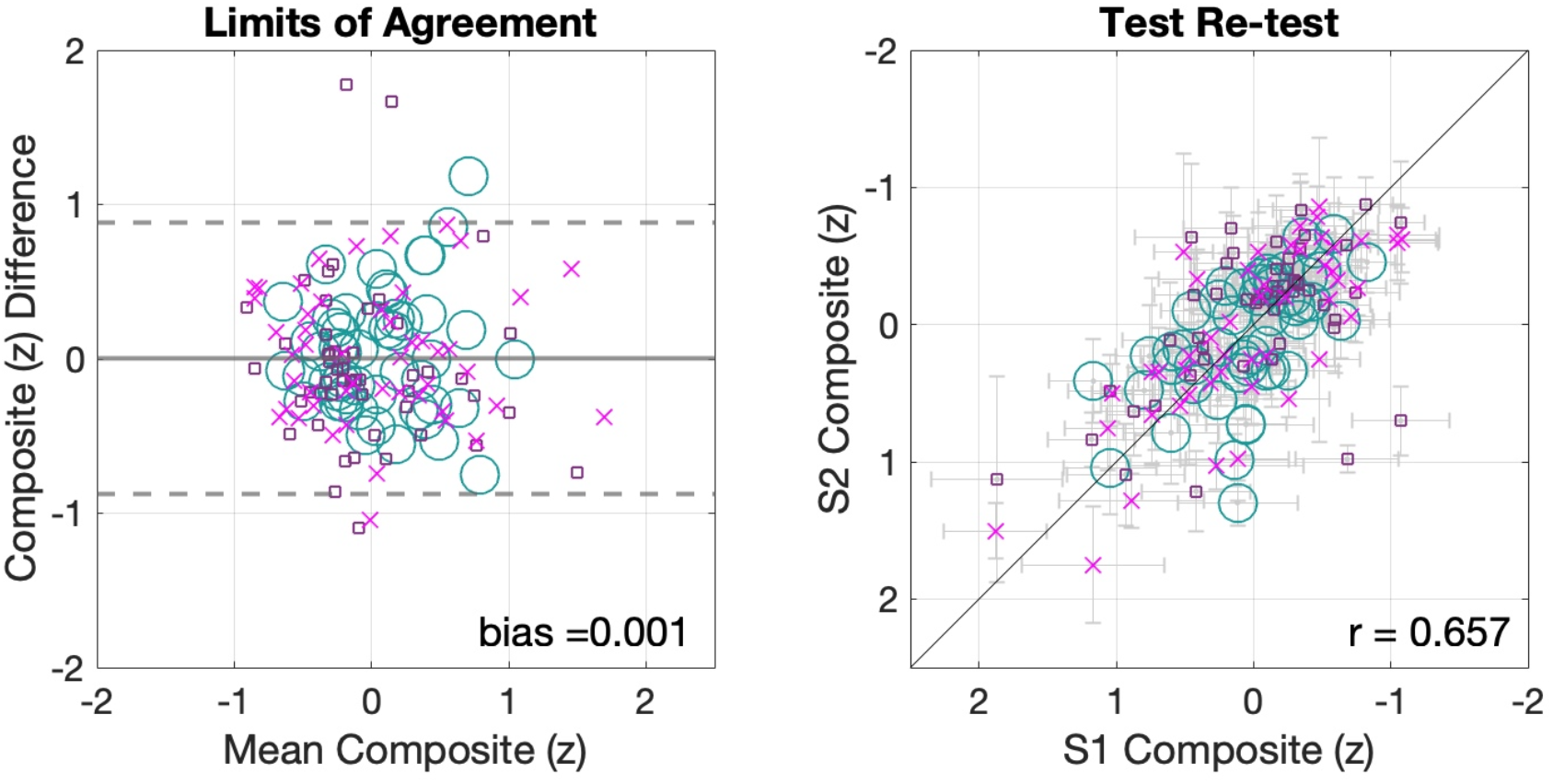
Composite Scores across all three conditions. Panel on the left shows the limits of agreement (see Altman & Bland, 1983) for the composite scores. Panel on the right shows scatterplot of composite scores. The horizontal error bars indicate SEM for session 1 while the vertical bars reflect session 2 SEM.

#### 1. Threshold Differences across Conditions

To address how composite scores changed as a function of listening condition, the composite score is plotted separately for each condition (Fig. 6). The Repeatability condition consisted of 51 participants who were tested in a quiet room and received the Sennheiser headphones with 30 dB passive attenuation that the system was calibrated on. The Headphone condition was similar to the Repeatability condition above except participants received either the Sennheiser 280 Pro headphones or the active-noise cancelling Bose Quiet Comfort 35 headphones. Finally, the Noise condition added environmental noise playing in the background at 70 dB on average and is otherwise identical to the methods reported in the Headphone condition.

**Fig 6.**
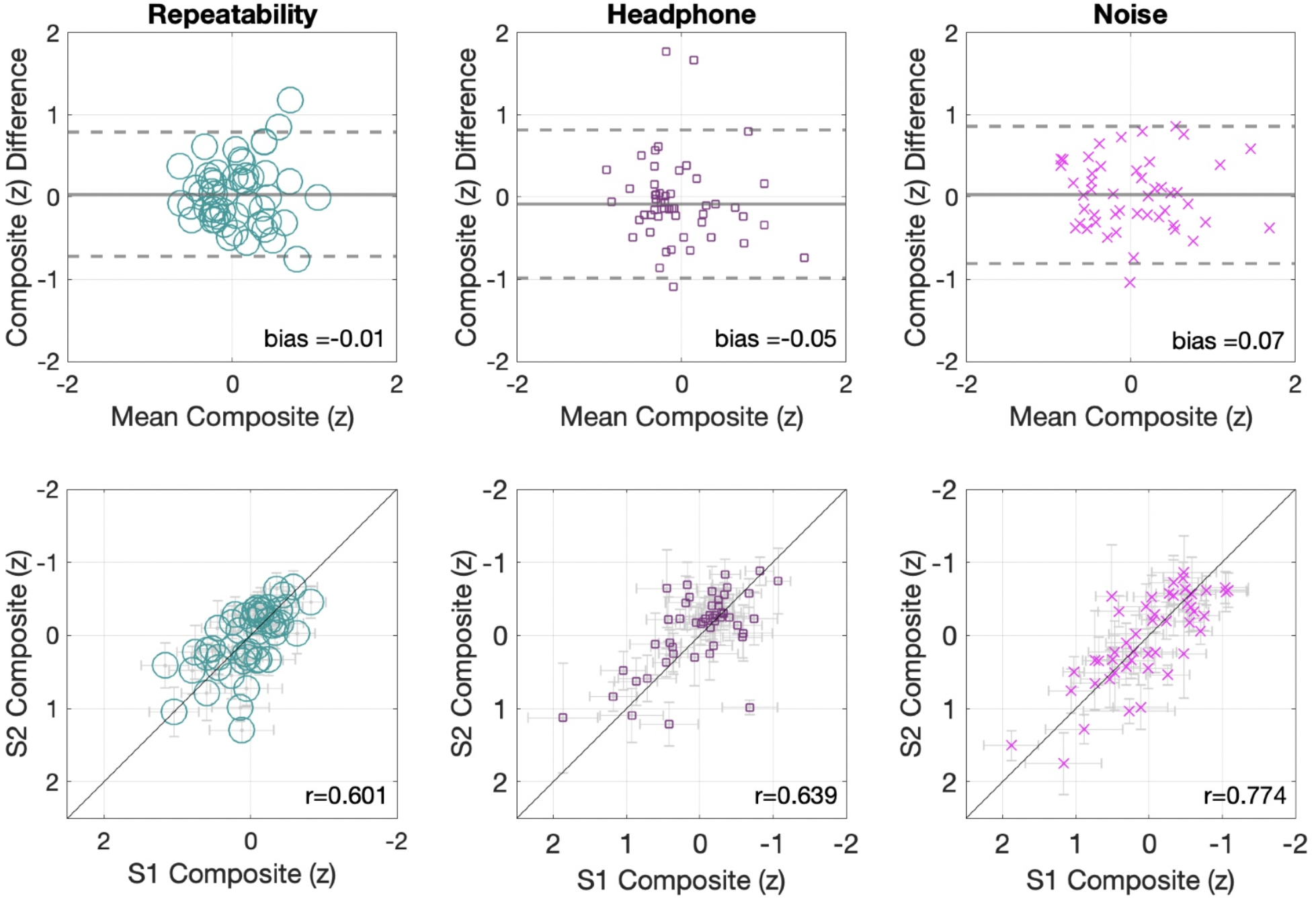
Composite scores for each condition. Top panels show the limits of agreement plots. Bottom panels show the composite scatterplots for each condition. The horizontal error bars indicate SEM for session 1 while the vertical bars reflect session 2 SEM.

In all three experimental conditions, composite scores showed minimal bias (Repeatability condition = −0.01; Headphone condition = −0.05; Noise condition = 0.07), limits of agreement that resemble the aggregate sample’s composite around 1 *SD* (Repeatability condition [−0.72, 0.79]; Headphone condition [−0.98, 0.81]; Noise condition [−0.8, 0.86]), and similar strength of association between scores of session 1 and 2 with (*r =* .601, *p* < .001) for Repeatability condition (standard); (*r =* .639, *p* < .001) for Headphone condition (silence); and (*r =* .774 *p* < .001) for Noise condition (noise). These correlations are within the 95% confidence intervals of the general aggregate composite *r*value.

To formally test the hypothesis of zero bias in threshold estimation between sessions for the different listening conditions, a series of t-tests were conducted. These tests failed to find significant differences in any of the conditions (Repeatability condition, *t_(50)_* = −0.34, *p =* .73, *Cohen’s d =* −0.04; Headphone condition, *t_(50)_* = 0.3, *p =* .76, *Cohen’s d* = 0.04); Noise condition, *t_(47)_* = −0.17, *p =* .86, *Cohen’s d =* −0.02). Finally, as an additional test of significance, a 3X2 repeated measures ANOVA with the within-subject factor Session and the between-subjects factor Condition was conducted to assess the overall effects of repeated measurements across a range of conditions. Again, no statistically significant effects were found for either Session (*F(_1,147)_* = 0.004, *p* = .94 *η^2^* < 0.01) nor for Condition (*F_(2,147)_* = 0.92, *p* = .4, *η^2^* = 0.01), and with no significant interaction (*F_(2,147)_* = 0.11, *p* = .88, *η^2^* < 0.01).

#### 2. Headphone Comparison

To examine the effects of headphone type and the presence of environmental noise, data are presented from the Headphone condition (both headphone types in silence) and the Noise condition (two headphone types in noise). Fig. 7 shows the limits of agreement between sessions as well as the scatter plots that show their association for the Headphone and Noise conditions separately. The data are plotted for each assessment in Supplemental Figures S2 and S3. The agreement analysis between the estimated thresholds using either set of headphones again showed minimally biased estimates (Headphone condition (silence) 0.003; Noise condition (noise) 0.082) and similar limits of agreement near 1 *SD* (Headphone condition (silence) [−0.94, 0.94]; Noise condition (noise) [−0.67, 0.84]) as reported in the general aggregate. Composite correlations were also similar to what is reported above, with the correlation for Headphone condition (silence) (*r =* .509 *p* < .001) suffering due to a reduced between-subject variability. A stronger association between measures was found in Noise condition (noise) where performance between-subjects is increased in relation to the within-subject variation (*r* = .74, *p* < .001).

**Fig 7.**
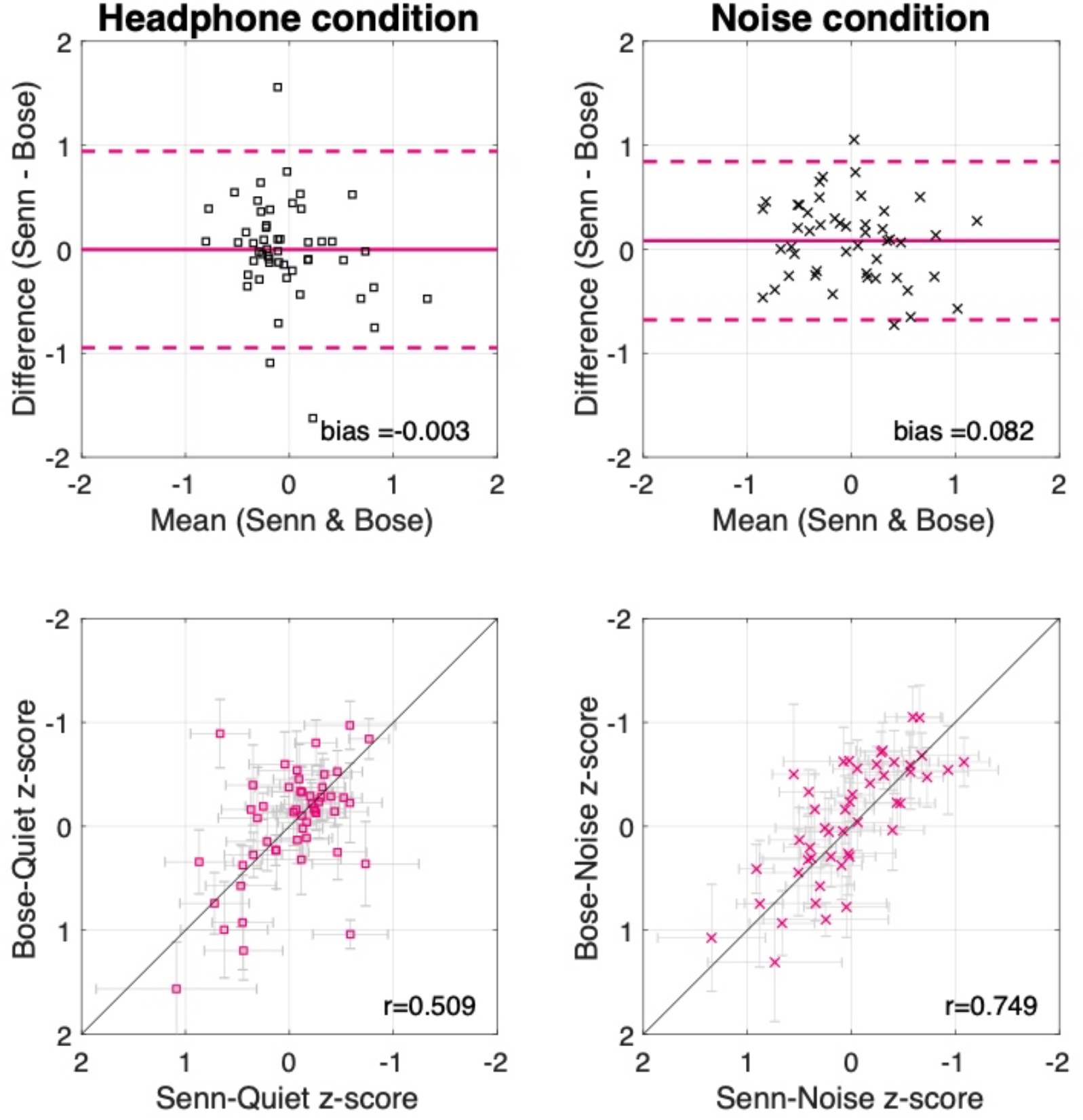
Headphone comparison. Top panels show the limits of agreement plots. Bottom panels show the composite scatterplots relating headphone type used. The horizontal error bars indicate Sennheiser 280 pro headphones.

The stability of threshold estimates across headphones in different environmental noise conditions is a notable result, as not only were the headphones different, but also, they shared the same output from the iPad, which was calibrated according to the Sennheiser 280 Pro—and not the Bose—headphones mechanical output levels as detailed in the Methods section. After calibration, an output level of 80 dB SPL (using the Sennheiser 280 Pro as recorded with a Brüel & Kjær Head and Torso Simulator with Artificial Ears in a VA RR&D NCRAR anechoic chamber) resulted in a level of 66 dB SPL for the Bose Quiet Comfort 35, with the high noise-cancelling setting engaged as used in all testing sessions (73 dB SPL with the noise-cancelling setting turned off). In order to allow the headphone effects to be examined without modification, and to avoid recalibration of the iPad between test sessions in the experiment, the settings that produced an 80 dB SPL output for the Sennheiser were used also for the Bose headphones. This meant that even in a silent environment, all of the stimuli were attenuated by 14 dB when Bose headphones were used.

Table 4 shows the mean thresholds and SDs for each type of headphone in each condition and assessment. Table 5 shows the within-subject LoA, correlations between headphones used, and repeated measures t-tests that formally test differences between the estimated thresholds with each headphone for each condition and assessment separately. The data associated with these statistical tests are plotted in Supplemental Figures S2 and S3. To test for differences in threshold estimation as a function of headphone type, t-tests were used to compare between the headphone types in each condition. Of note, since headphone type was counterbalanced across sessions, these analyses were averaged across sessions. No statistically significant effects were observed in either condition (Headphone condition (silence), *t_(50)_* = −0.03, *p* = .97, *Cohen ’s d* = −0.005; Noise condition (noise) *t_(47)_* = 1.45, *p* = .15, *Cohen ’s d* = 0.21). As an additional test of significance, a 2X2 repeated measures ANOVA with the within-subject factor Headphone and the between-subjects factor Condition was conducted to assess headphone effects. Again, no statistically significant effects were found for either Headphone (*F_(1,97)_* = 0.8, *p* = .37 *η^2^* < 0.01) nor for Condition (*F_(1,97)_*= 0.01, *p* = .91, *η^2^* < 0.01), and with no significant interaction (*F_(1,97)_* = 0.9, *p* = .34, *η^2^* < 0.01).

**Table 4.**
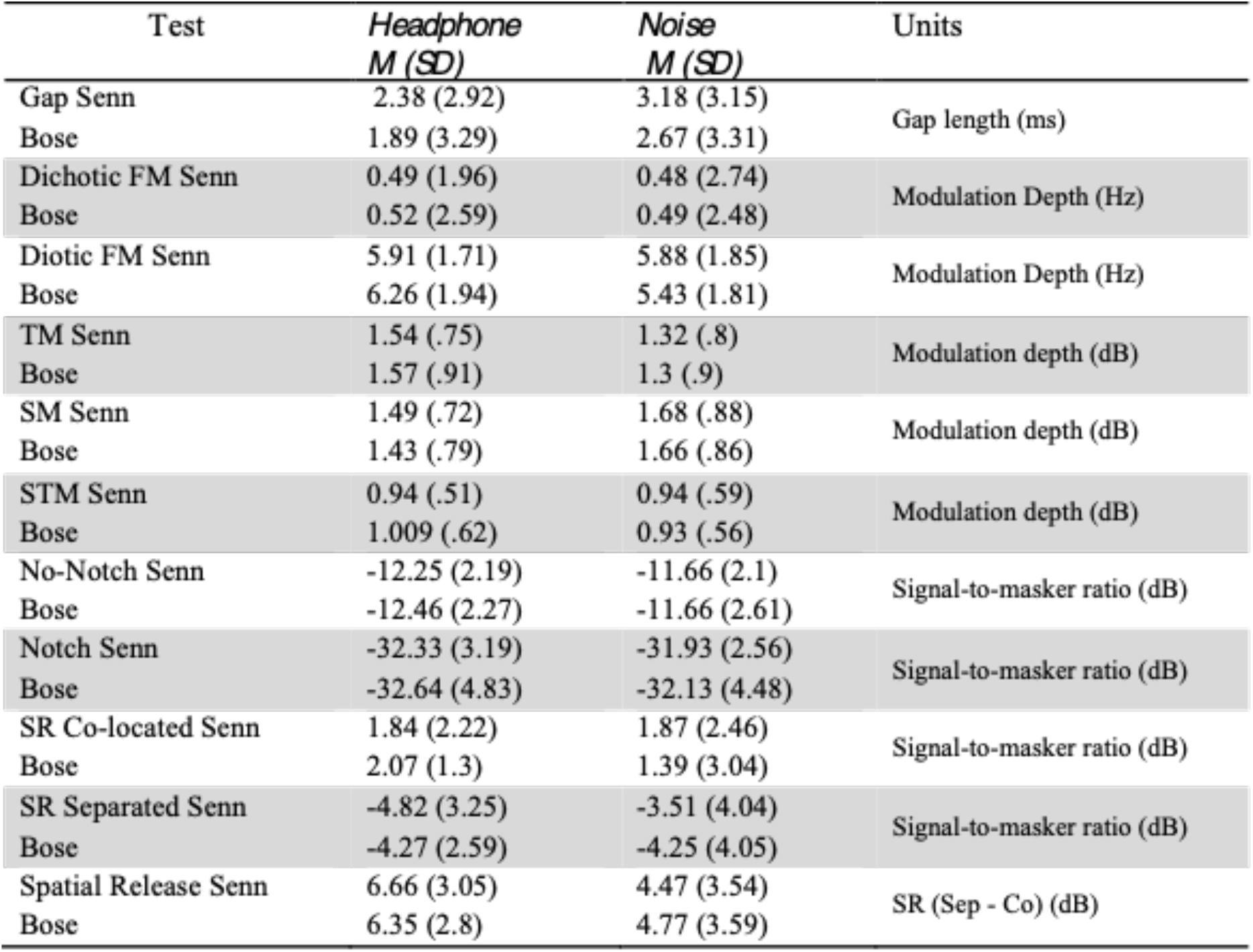
Mean thresholds and standard deviations for the 10 assessments utilized plus the derived spatial release metric across both conditions that used different headphones. Data is presented in PART’s native measurement units except for the targets-in-competition tests that have been converted to TMR. The first row of each test shows thresholds obtained with the Sennheiser 280 Pro system and the second with the Bose Quiet Comfort 35 system.

| Test | Headphone<br>*M (SD)* | Noise<br>*M (SD)* | Units |
| --- | --- | --- | --- |
| Gap Senn | 2.38 (2.92) | 3.18 (3.15) | Gap length (ms) |
| Bose | 1.89 (3.29) | 2.67 (3.31) |  |
| Dichotic FM Senn | 0.49 (1.96) | 0.48 (2.74) | Modulation Depth (Hz) |
| Bose | 0.52 (2.59) | 0.49 (2.48) |  |
| Diotic FM Senn | 5.91 (1.71) | 5.88 (1.85) | Modulation Depth (Hz) |
| Bose | 6.26 (1.94) | 5.43 (1.81) |  |
| TM Senn | 1.54 (.75) | 1.32 (.8) | Modulation depth (dB) |
| Bose | 1.57 (.91) | 1.3 (.9) |  |
| SM Senn | 1.49 (.72) | 1.68 (.88) | Modulation depth (dB) |
| Bose | 1.43 (.79) | 1.66 (.86) |  |
| STM Senn | 0.94 (.51) | 0.94 (.59) | Modulation depth (dB) |
| Bose | 1.009 (.62) | 0.93 (.56) |  |
| No-Notch Senn | -12.25 (2.19) | -11.66 (2.1) | Signal-to-masker ratio (dB) |
| Bose | -12.46 (2.27) | -11.66 (2.61) |  |
| Notch Senn | -32.33 (3.19) | -31.93 (2.56) | Signal-to-masker ratio (dB) |
| Bose | -32.64 (4.83) | -32.13 (4.48) |  |
| SR Co-located Senn | 1.84 (2.22) | 1.87 (2.46) | Signal-to-masker ratio (dB) |
| Bose | 2.07 (1.3) | 1.39 (3.04) |  |
| SR Separated Senn | -4.82 (3.25) | -3.51 (4.04) | Signal-to-masker ratio (dB) |
| Bose | -4.27 (2.59) | -4.25 (4.05) |  |
| Spatial Release Senn | 6.66 (3.05) | 4.47 (3.54) | SR (Sep - Co) (dB) |
| Bose | 6.35 (2.8) | 4.77 (3.59) |  |

**Table 5.** Limits of agreement and significance testing for the 10 assessments comparing headphones used in two conditions. The first row shows the Headphone condition and the second the Noise condition. Positive values on the bias column indicate better performance with the Sennheiser system except on the Spatial Release metric which is the only scale in which bigger magnitudes indicate better performance. * indicate significance at α = .05

| Test | Bias | Limits of Agreement | Units | $r(p)$ | $t(p)$ | Cohen's $d$ | df |
| --- | --- | --- | --- | --- | --- | --- | --- |
| Gap | -0.33 | [-3.63 to 2.97] | Log2 | .47 (<.01)* | 1.37 (.17) | -.2 | 48 |
| (Noise) | -.25 | [-3.36 to 1.67] | (ms) | .56 (<.01)* | 1.07 (.28) | -.14 | 45 |
| DichoticFM | 0.07 | [-2.62 to 2.77] | Log2 | .35 (.01)* | -0.37 (.7) | .06 | 50 |
| (Noise) | 0.02 | [-2.21 to 2.25] | (Hz) | .66 (<.01)* | -0.12 (.9) | .01 | 47 |
| DioticFM | 0.08 | [-1.88 to 2.05] | Log2 | .34 (.01)* | -0.58 (.56) | .96 | 47 |
| (Noise) | -0.11 | [-1.72 to 1.49] | (Hz) | .56 (<.01)* | 0.94 (.34) | -.12 | 46 |
| TM | 0.03 | [-1.58 to 1.64] | M | .53 (<.01)* | -0.28 (.77) | .03 | 50 |
| (Noise) | -0.01 | [-1.7 to 1.67] | (dB) | .49 (<.01)* | 0.13 (.89) | -.02 | 44 |
| SM | -0.06 | [-1.55 to 1.43] | M | .5 (<.01)* | 0.55 (.58) | -.08 | 46 |
| (Noise) | -0.01 | [-1.78 to 1.75] | (dB) | .47 (<.01)* | 0.11 (.9) | -.01 | 43 |
| STM | 0.06 | [-1.19 to 1.32] | M | .38 (<.01)* | -0.71 (.47) | .11 | 47 |
| (Noise) | -0.004 | [-1.45 to 1.45] | (dB) | .19 (.23) | 0.03 (.97) | -.007 | 39 |
| No-Notch | -0.2 | [-5.02, 4.6] | SMR | .39 (<.01)* | 0.59 (.55) | -.09 | 49 |
| (Noise) | 0 | [-6.1 to 6.1] | (dB) | .14 (.34) | 0 (1) | 0 | 43 |
| Notch | -0.31 | [-8.72 to 8.09] | SMR | .49 (<.01)* | 0.51 (.6) | -.07 | 49 |
| (Noise) | -0.2 | [-9.52 to 9.11] | (dB) | .17 (.25) | 0.27 (.78) | -.05 | 41 |
| Co-located | 0.23 | [-3.77 to 4.24] | SMR | .42 (<.01)* | -0.82 (.41) | .12 | 50 |
| (Noise) | -.48 | [-5.89 to 4.91] | (dB) | .51 (<.01)* | 1.13 (.26) | -.17 | 40 |
| Separated | 0.54 | [-6.31 to 7.4] | SMR | .29 (.03)* | -1.12 (.26) | .18 | 50 |
| (Noise) | -.74 | [-6.32 to 4.83] | (dB) | .75 (<.01)* | 1.79 (.07) | -.18 | 46 |
| SpatialR | -0.31 | [-8 to 7.37] | SMR | .107 (.45) | 0.57 (.57) | -.1 | 50 |
| (Noise) | 0.3 | [-6.28 to 6.88] | (dB) | .55 (<.01)* | -0.56 (.57) | .08 | 39 |

In summary, the data failed to show any systematic effect of headphone type when participants were tested in either silent or noisy environments. These composite analyses further support the reliability of PART and suggest that it may be achieved with or without active noise cancelling technology and in presence of moderate environmental noise. These results also suggest that even a 14-dB difference at the mechanical output level did not produce noticeable differences in performances for these undergraduate students with hearing in the normal range.

### E. Effects of Staircase Parameters

A final difference present between Conditions was the ratio of step-sizes up:down used in the adaptive staircases. This was 2:1 for the Repeatability Condition and 1.5:1 for the other two (Headphone & Noise). This modification was inspired by the work of García-Pérez (2011) and the implications are still being explored by work such as Hoover et al. (2019). This modification was not applied to the spatial release tasks, for which the progressive structure remained unchanged. The results revealed that this manipulation did not manifest salient differences in terms of the estimated thresholds between conditions, however it did have an impact on the number of trials required to achieve a threshold estimate. This is shown in Figure 8, in which the mean number of trials are plotted per task per experiment.

**Fig 8.**
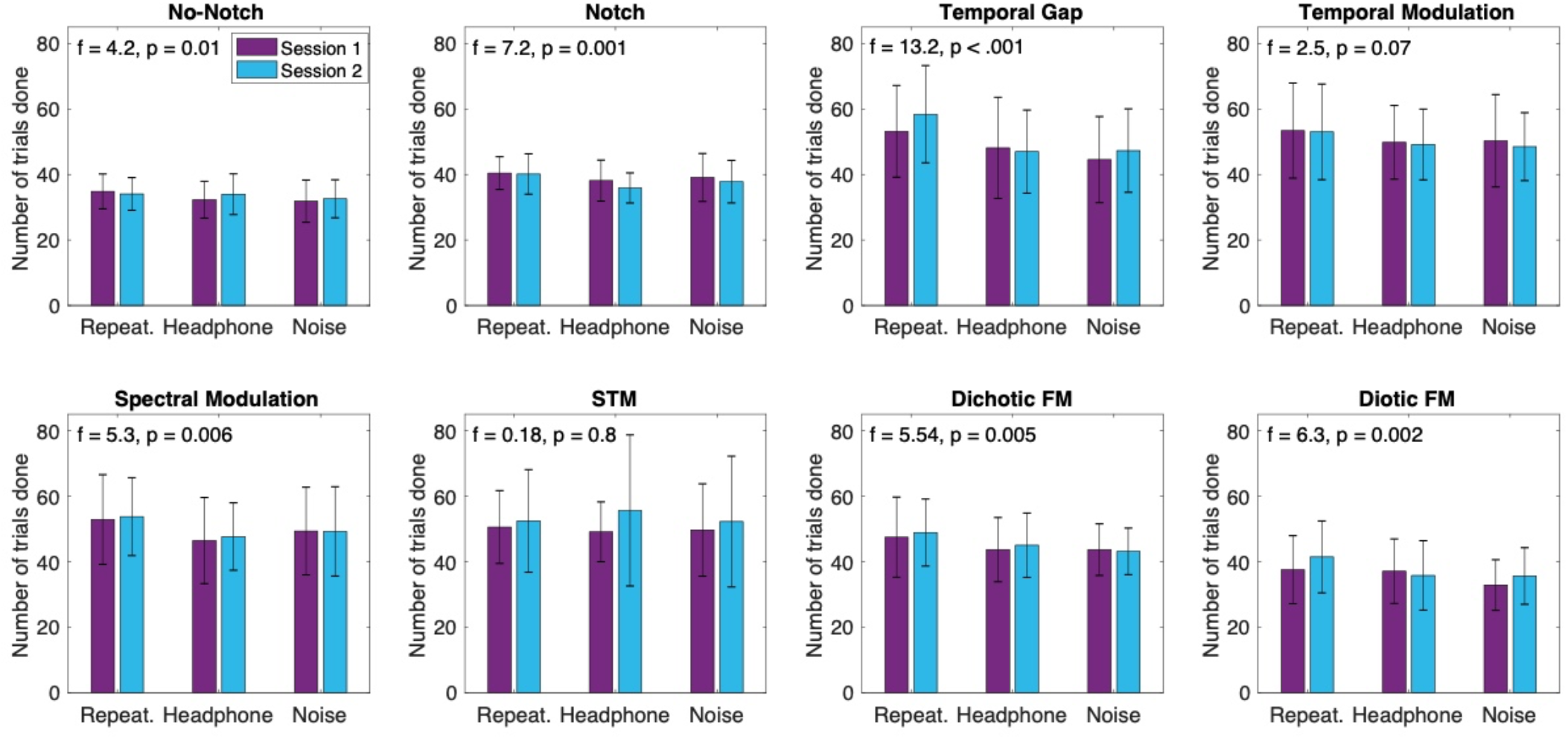
Mean and standard deviations of the number of trials presented per task for each Experiment. Statistics from a one-way ANOVA with the between-subject factor Condition (3 levels) are displayed in the top of each graph.

To address the effects specific to the staircase, a series of independent samples t-tests were conducted between the number of trials needed to achieve a threshold estimate in the Repeatability vs Headphone conditions. These two conditions are the most similar (apart from the headphone differences in one of the sessions of the Headphone condition) and so are the best place for examining the difference between step size ratios. It would have been possible to conduct this comparison only on the conditions using the same headphones, but this would have reduced the size of the data set in half. The number of trials for the adaptive staircases are included in the Supplemental dataset to permit alternative analyses to be conducted. Differences that met statistical significance were obtained in a number of the assessments, supporting the hypothesis that tests with the 1.5:1 staircase were more likely to finish in fewer trials. The effect sizes were relatively small, as differences are no more than 6 trials on average and sum up to 25.8 trials on average for the whole battery. On the other hand, there were no significant differences between the number of trials for any of the tests of the Headphone and Noise conditions, both of which used the 1.5:1 ratio. These results indicate the change in step-size from 2:1 to 1.5:1 resulted in staircases that were slightly more efficient but equally reliable in threshold estimates. It is, of course, the case that every study to which data were compared above, and most in the literature, used equal step-sizes (Levitt, 1971), so further research will be required to determine whether there is an advantage, or cost, related to using the uneven step-sizes chosen in the current study. Because the results are highly similar across all three experiments (see Fig. 3), and our main intention with this dataset is to provide normative threshold estimates of central auditory function in a variety of settings to potentially supplement clinical practice, the results are described a manner that focuses on the combined data set across experiments. This analysis uses composite measures to address the differences between experiments (aggregating across tests). More detailed reports about the tests in each experiment can be found in the supplemental materials.

## IV. Discussion

This study examined the validity and reliability of a battery of ten assessments that evaluate different aspects of central auditory function using the Portable Automatic Rapid Testing (PART) application applied to young adult listeners without reported hearing problems. Overall, results show that thresholds can be obtained that are highly consistent across sessions and that are very similar to those reported in laboratory settings obtained with more traditional equipment and, in some cases, with extended testing and training (see Table 3). Furthermore, results from the Repeatability condition were replicated in the Headphone and Noise conditions, demonstrating that PART produces consistent threshold estimates across a variety of settings and equipment. Overall these results suggest that the PART platform can provide valid measurements across a range of listening conditions.

An important utility for this study is that it provides an initial normative data set for young adult listeners for the PART tasks reported, and eventually as a reference for patient populations. However, substantial work is required before PART will be appropriate for clinical use. For example, while the tasks included in this first battery were chosen based upon prior literature suggesting possible sensitivity in understanding listening disorders, these data do not capture variations in age and do not include effects of differences in hearing threshold (see Jakien and Gallun, 2018). Future work will involve developing a similar normative data set for this battery. With such a dataset it would be possible to determine which combinations of tests are most sensitive in distinguishing between different disorders. Likewise, the reliability of measures needs to be established in different populations that may have more difficulty with the procedures than the college student population reported here. The number of outliers observed in the current data set (see below for further discussion) provide a lower bound on the number of expected false-tests. Further, although learning effects are among the smallest effects observed in this data set, they must be explored in relevant patient populations and potentially accounted for when interpreting test results. In both cases, our results suggest that and either repeated measures or adjustments to adaptive procedures will be required to increase reliability in patient populations. Still, the fact that thresholds similar to those found in the literature can be obtained on a large number of tests within a short-period of time, using consumer-grade technology, provides optimism that PART will be useful in the clinic.

The criteria we used for outlier rejection was justified by the goal of creating a normative dataset, but it is important to note that the field holds a variety of different views regarding outlier rejection. The current choice is simply definitional, in that ‘normative’ refers to a normal distribution, and thus, it is appropriate to reject data falling far outside of this distribution. Nevertheless, there was a minimal effect on population estimates of means and standard deviations, whether or not outliers are excluded (see Supplemental Table 1). Supplemental Figure 1, which shows the data with outliers circled, reveals that the main impact of including outliers is to make it more difficult to see the normal range of the dataset. Another important question is whether or not it is possible to say something meaningful about which listeners gave data that was then rejected. For example, are they impaired in auditory processing, or do they represent typical variation of the larger population? It seems unlikely that these participants had any severe hearing loss as they all self-reported to have no hearing difficulties, which is considered a reasonable indication for normal hearing (Vermiglio, Soli & Fang, 2017), and were able to easily detect a 45 dB 2kHz pure tone which assured an audibility minimum criteria. Moreover, as indicated in Supplemental Table 1, most outlying cases were not consistent across sessions, and as can be seen in Supplemental Figure S1 are within the normal range in one of the testing sessions. This suggests that many of the outliers were either inattentive, unmotivated or confused or otherwise non-compliant in one of the sessions. Further some of the outliers were actually in the supra-normal hearing range, again evidence against outliers being indicative of hearing impairment. Still, while it is reasonable to suggest that outliers do not represent normative or dominantly systematic effects, they are still a concern and do need to be considered when contemplating clinical implications, especially of a single test. In particular, it is important to keep in mind the expected probability (ranging from 1-8% depending on the test) of getting an unreliable test result when using these tests on this platform.

The adaptive procedures employed, may have also contributed to extreme outlying data points; they can be particularly important when evaluating individual test reliability. It is reported above that changes to the parameters of the adaptive procedure can impact the efficiency of estimating a threshold; however, it is also likely that changes to the procedure itself can add to reliability. Of note, while 3-down 1-up staircases are typical in laboratory settings, it is common to observe outlying tracks, especially in 2-AFC tasks that have a 50% guessing rate. However, instead of simply averaging across reversals, as is conventional, one can also look at the standard deviation across the reversals to determine the extent to which the staircase is converging consistently. In cases of high variance across reversals, or also in cases where data are far outside of the norms, it may be prudent to either repeat staircase runs and/or repeat procedures. Likewise in the case of the Spatial Release from Masking Task, differences between progressive tracks can be used to determine whether additional tracks should be run. PART has implemented a number of adaptive procedures and these and other reliability checks can easily be implemented and evaluated when addressing individual test reliability, as opposed to population test reliability as evidenced here.

Another issue of concern is the extent to which performance may change systematically across testing sessions. LoA plots, correlational analyses, and statistical tests of session effects all provide complementary information on changes in performance over time. In general, systematic bias of threshold estimates across sessions was minimal, and similar to the bias observed across levels of performance (see LoA plots). The expected differences among measures across sessions are estimated in the limits of agreement between sessions. While significance testing revealed some differences between sessions in some of the tests, the effect sizes we found are small and typically comparable to the smallest step sizes used in the procedures. Further, these can now be considered as test re-test effects in future work. Consistent with this, reliability was further quantified using Pearson r, which is not sensitive to systematic effects of testing session. Although correlation is highly reactive to small changes in between-subject variability (see Supplementary Table 1), and cases with reduced between-subject variability, it presents complimentary information regarding the relation among within-subject and between-subject variabilities. When the assessments with smaller r values in complement to the LoA plots are examined, it can be seen that in most cases the limits of agreement closely resemble other tests with higher r values. This is an indication that r is decreasing due to restricted ranges of good performance as was to be expected in this sample of normal listeners.

It is notable that thresholds were consistent across different external noise conditions (Repeatability & Headphone vs Noise conditions) and/or used different types of headphones (Repeatability vs Headphone & Noise conditions) (see Figs. 7 & 8). Also of note, the correlations were higher for the condition with external cafeteria noise without an increase on the limits of agreement. This is important because a test platform that is portable, automatic, and rapid can only be successfully exploited if it is able to provide accurate measurements that can be collected in a variety of potentially less than optimal settings. Here, we have shown that PART was able to obtain estimates of central auditory function that resemble those found under laboratory conditions, despite using untrained listeners tested in settings as noisy as a university cafeteria. PART should be considered as a supplemental tool in the clinic which can be used to collect valuable information about a person’s hearing capabilities with little need for supervision from a clinician. These results also suggest that this system and these tests are robust to the presence of moderate noise and substantial variability in sound output levels.

This study also compared the use of headphones with an active noise-cancelling technology to those with passive attenuation. We considered it worthwhile to test this technology because it is now widely available, but little is known about the advantages and disadvantages it could represent for auditory testing. The active noise cancelling algorithm that these headphones (Bose) use could potentially aid or distort the perception of the sounds used in our tests. We failed to find a statistically significant effect between threshold estimates obtained for the different headphones under both in silent and noisy listening conditions. In other words, estimated thresholds were similar for the Sennheiser 280 Pro in silence (Headphone condition), and this lack of difference manifested similarly in noisy conditions (Noise condition). This suggests that the passive attenuation provided by the Sennheiser 280 Pro is sufficient to obtain reliable measurements in less than optimal external noise conditions outside of the sound-booth. Also, it suggests that the differences between the headphones, including the active noise-cancelling algorithm, are not changing the signal in any way that results in significant reductions in performance. Perhaps the noise-cancelling signal processing was inactive or operating at low frequencies that did not affect performance. In any case, threshold estimation held constant across the headphone technologies used with a single calibration profile (the same output from the iPad). These data serve as verification that relatively inexpensive auditory hardware can be used to test auditory function in a variety of settings with sufficient precision to provide clinical evidence of central auditory function in individual listeners.

PART can thus appropriately be considered as a valid platform for testing central auditory processing. It is robust to moderate levels of ambient noise and small variants in equipment and procedure. The reported data can now be used as a normative baseline against which auditory dysfunction can be identified in future work. However, clinical research will be needed to determine how thresholds vary as a function of age and different degrees of hearing loss. The reliability analysis reported here applies only to young listeners with normal hearing, future work will need to address whether threshold estimates from PART can be reliably obtained for older listeners with varying degrees of hearing loss, and to determine the extent to which the measured reliability in this work is adequate for identifying central auditory processing deficit. This next step is feasible considering that the PART platform is highly accessible given its relatively low cost in expense (it only requires a computer tablet and headphones), time (the whole battery of 10 assessments in under 1 hr), human resources (it runs the assessments automatically, one after another, including instructions and breaks), and that it can be used in range of environmental settings suitable for testing (from the anechoic chamber as in Gallun et al. (2018) to noisy cafeteria conditions). Thus, PART has the potential to provide a supplementary tool to gather the quantity and variety of psychophysical measures of auditory function that will allow us to translate laboratory findings into the clinic to inform clinical practice.

## Acknowledgment

We are grateful to Michelle Molis, Kasey Jakien and Sittiprapa Isarangura for their kind revisions to the manuscript. This work was funded by NIH NIDCD R01 DC 015051. Equipment and engineering support provided by the VA RR&D NCRAR, the UCR Brain Game Center, and Samuel Gordon (NCRAR). The first author is currently funded by CONACYT and UC Mexus. The views expressed are those of the authors and do not represent the views of the NIH, CONACYT, UC Mexus, or the Department of Veterans Affairs.

