## Supplementals for "Portable Automated Rapid Testing (PART) for auditory assessment: Validation in a young adult normal-hearing population"

### Supplemental Materials

#### ***Outlier Analysis***

Table ST1 is provided to demonstrate that outlier rejection, which was largely for ease of presentation has a limited impact on estimates of normative ranges for the assessments (compare columns 4 to 5 of table ST1). Figure S1 plots the data from all participants for all tests, with outliers indicated as circled symbols. Table ST1 also provides measures of how many outliers were observed in each condition of each task and the degree to which a given participant provided consistent thresholds, using a metric of a score greater than 2 standard deviations beyond the mean on both sessions, as shown in columns 2 and 3 in Table ST1. This reveals that for those participants for whom data were excluded, thresholds were typically within the normal range in one of the sessions. This is also visible in Figure S1.

**Table ST1.** Outlier cases, consistency across sessions, and impact on mean thresholds and standard deviations for the 10 assessments and the spatial release metric.

| Assessment | Outliers (by condition) | Consistent cases <\|2SD\| ΔSession | M (SD) Full-data | M (SD) No-outliers | Units |
|---|---|---|---|---|---|
| Gap | 4 (0, 2, 2) | 0 | 2.49 (2.9) | 2.36 (2.75) | ms |
| DichoticFM | 2 (2, 0, 0) | 2 (2, 0, 0) | 0.53 (2.1) | 0.52 (2) | Hz |
| DioticFM | 5 (1, 3, 1) | 2 (0, 2, 0) | 6.3 (1.7) | 6.1 (1.6) | Hz |
| TM | 4 (1, 0, 3) | 2 (0, 0, 2) | 1.59 (.93) | 1.49 (.73) | (M) dB |
| SM | 9 (1, 4, 4) | 4 (0, 1, 3) | 1.71 (.99) | 1.52 (.63) | (M) dB |
| STM | 13 (2, 3, 8) | 4 (0, 1, 3) | 1.18 (.86) | 0.95 (.4) | (M) dB |
| No-Notch | 5 (1, 0, 4) | 1 (1, 0, 0) | -11.6 (2) | -11.8 (1.7) | SMR dB |
| Notch | 7 (0, 1, 6) | 0 | -30.9 (6.1) | -32 (2.9) | SMR dB |
| SR Co-located | 7 (0, 0, 7) | 5 (0, 0, 5) | 1.51 (2.5) | 1.94 (1.6) | SMR dB |
| SR Separated | 2 (1, 0, 1) | 0 | -4.34 (3.1) | -4.47 (2.9) | SMR dB |
| Spatial Release | 9 (1, 0, 8) | 5 (0, 0, 5) | 5.86 (3) | 6.19 (2.79) | dB |

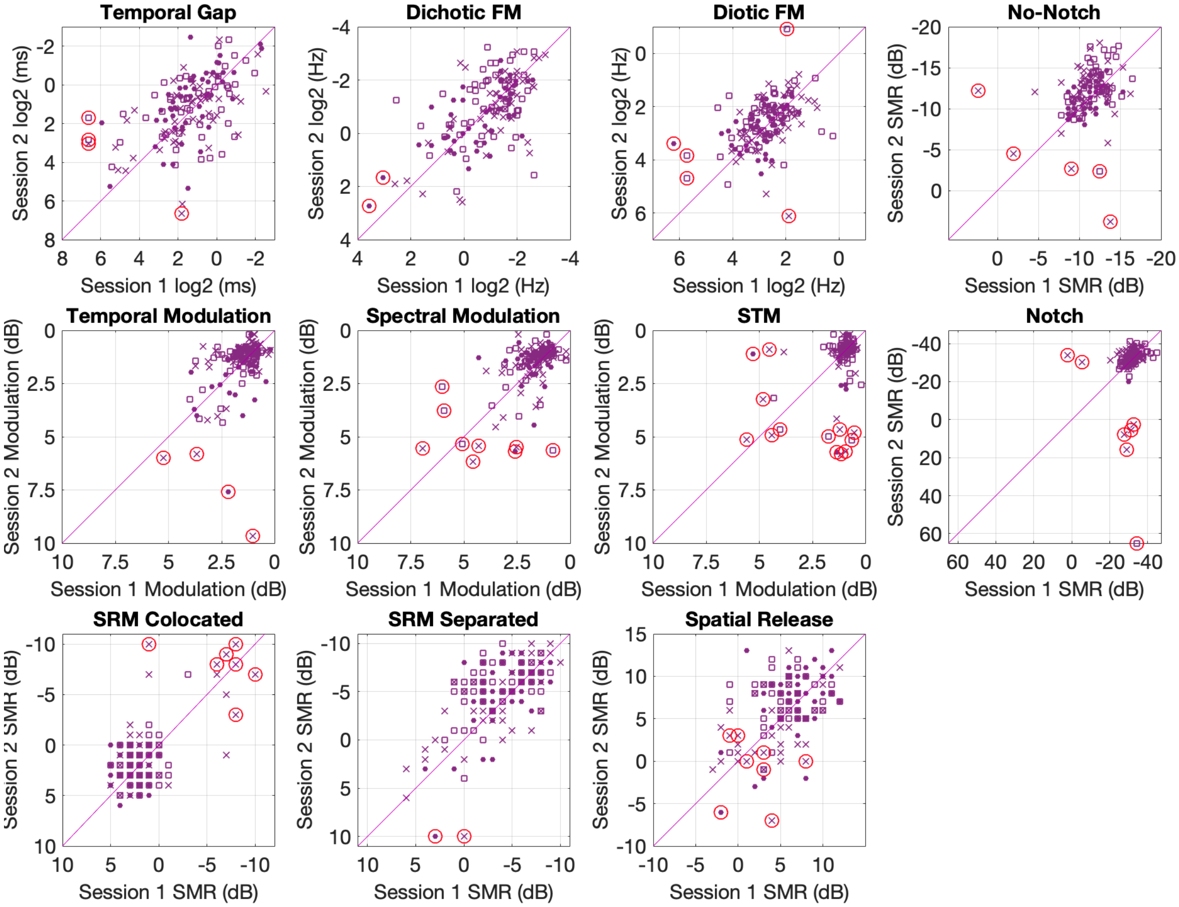


**Figure S1**. Scatter plots of Session 1 vs Session 2 for the 10 assessments used for all three conditions. Filled circles represent the Repeatability condition, open squares represent the Headphone condition, and crosses represent the Noise condition. Cases flagged as outliers (±3 *SD*) and removed from main analysis are marked with a surrounding circles. All axes are oriented to show better performance values away from the origin. The diagonal is plotted to ease evaluation of differences between sessions, dots above this line indicate better performance in session 2.

### ***Headphone effects***

Figures S2 (Limits of Agreement) and S3 (scatterplots) show the within-subject effects of headphone type.

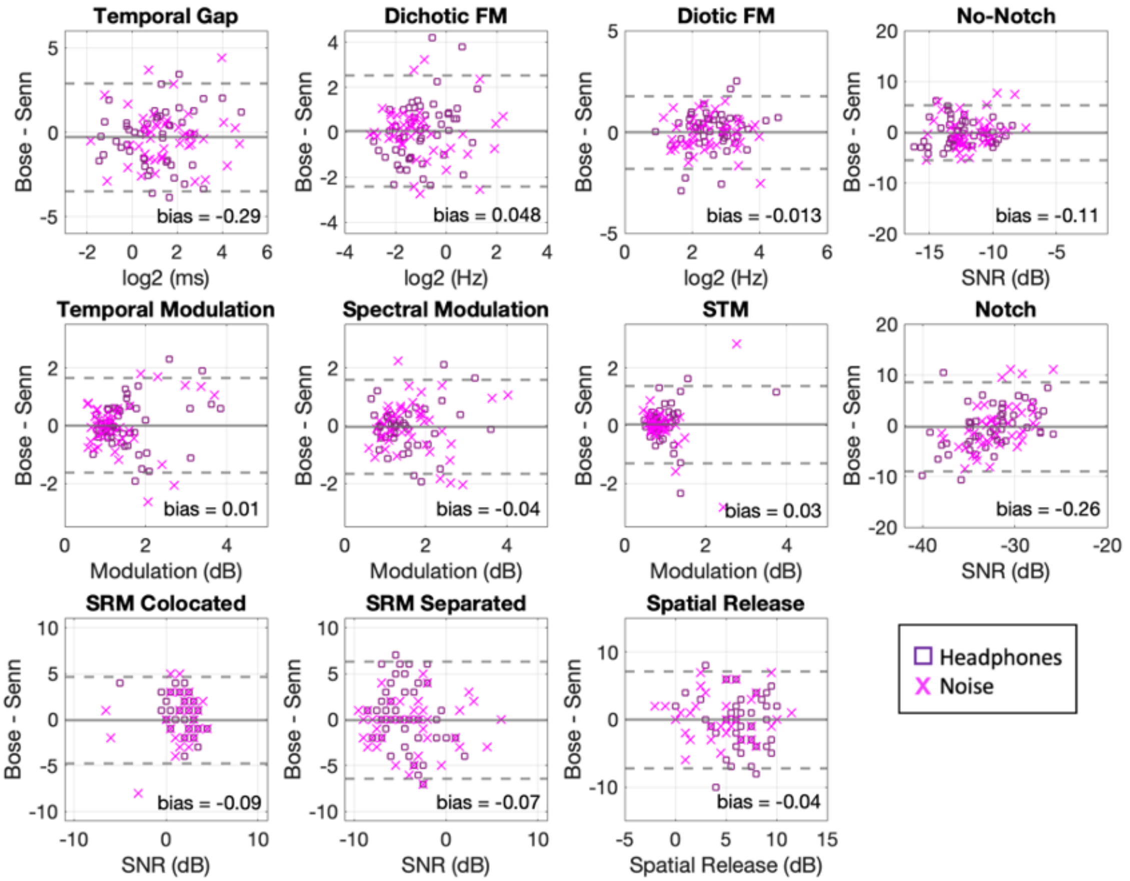


**Figure S2**. Limits of agreement of the estimated thresholds across headphone types in conditions 2 (squares) and 3 (crosses). The solid lines indicate the mean difference between headphone type. Dotted lines indicate the 95% limits of agreement. The red circle indicates the mean threshold for each test centered at zero difference between headphones. Solid lines below zero indicate better performance on the Bose headphones with active noise attenuation for all the plots except the spatial release metric, for which higher values indicate better performance.

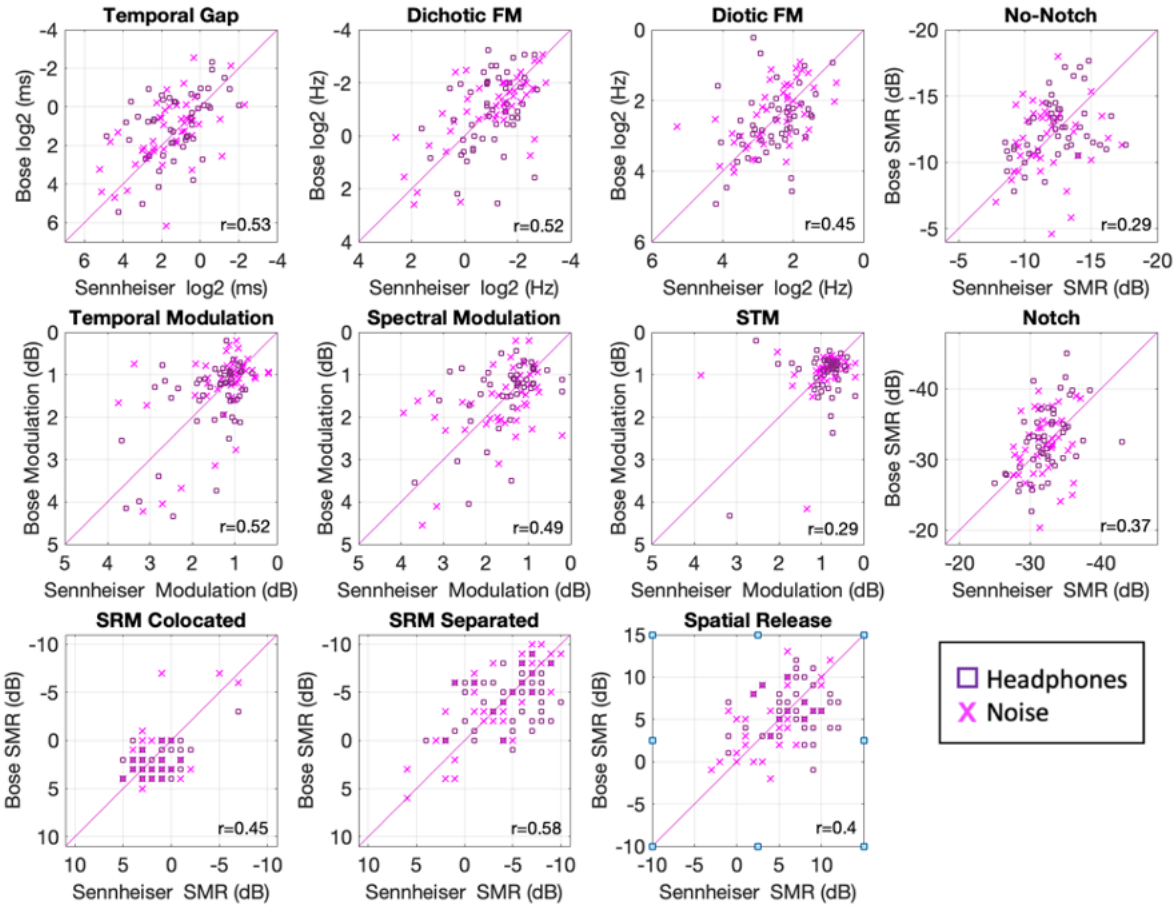


**Figure S3**. Scatter plots relating headphone types used for the 10 assessments in conditions 2 (squares) and 3 (crosses). All axes are oriented to show better performance values away from the origin. Correlations are indicated in the lower right of each panel. The diagonal is plotted to ease evaluation of differences between headphone types, dots above this line indicate better performance with active noise attenuation.

### ***Instructions***

Informed consent was obtained by a research assistant or experimenter before testing started. All participants were given demographic surveys and then heard the following instructions, read by a research assistant or experimenter:

> "We are developing a set of tests with diverse types of sounds that will help us better understand your hearing abilities. As we age (or by injury) our hearing abilities decline, and it is important to better understand the nature of this loss so we can deal with diagnosis and rehabilitation. During your session, which will take roughly 45 minutes, you will be listening to sounds and responding to them on an iPad. Your participation will aid research aimed to help improve people's quality of life, please take it seriously and give it your best effort."

Participants then were taken to the test room, where they were given an iPad and set of headphones. They were verbally instructed to sit down and put the headphones on with the left phone aligned with the left ear. All were then read the following instructions:

> "Now you will be tested on several different tasks. The instructions will be shown to you each time a task begins. The game is programed to challenge your hearing limits, so it will get harder to solve as you go. Please try to respond to the best of your ability, the better you are able to perform, the quicker the program will find your limit and the task will end. This experiment is divided in four chunks, now you will start the first one, when it finishes, the instructions will ask you to call me so I can write down your scores. Please follow the instructions on the iPad carefully. Do you have any questions?
>
> I will stay in here with you for the first few trials in case you have any further questions. Good luck!"

The first thing that appeared to them on the iPad was an instruction screen that read:

> “Welcome to PART! In this experiment, you will be responding to different series of sounds to test your hearing. In this first example, 4 squares will be presented, and will light up sequentially one after another. Your job is to find and touch the square that makes a sound as it lights up.”

These instructions were followed by a familiarization/screening task involving ten trials in which they were required to detect a 2kHz tone presented in one of the two test intervals, with silence presented during the other three intervals. Performance was monitored and instructions were repeated if necessary. Some participants did not anticipate that the noise would be near detection threshold and so did not hear anything. They were re-instructed, the testing was restarted, and then all were able to detect at least 9 of the ten targets. At this point, all participants completed conditions No-Notch and Notch from the Targets in Competition battery, before which the following instructions were shown on the iPad screen: ,

> “Next, you will be looking for the same bip sound, but this time, noise will play on every square. Your job is to find the square that makes a bip sound in addition to the noise. The program will try to find the amount of noise necessary for you to be unable to find the bip sound, just try your best and guess if uncertain. Hint: the first and last squares never contain the bip.”

Participants then were randomly assigned to complete the second half of the Signals in Competition Battery (SRM assessments), the STM battery or the TFS battery.

The STM battery used similar descriptions for all three assessments it contained. It started with the following instructions:

> “One of these sounds is not like the others... On every trial, four squares will be presented on screen. They will light up and emit a sound, one at a time. The first and the last squares will always carry some type of "ordinary" sound. One of the two squares in the middle will have a "special" modification, the other will carry "ordinary" sound. Your task is to identify which of the squares in the middle carries the "special" sound.”

The TFS battery contained the following instructions for Dichotic FM:

> "One of these sounds wobbles... On every trial, four squares will be presented on screen. One of the two squares in the middle will have a "special" modification, the sound they emit will seem to wobble between your ears! Your task is to identify which of the squares in the middle carries the "special" wobbling sound."

The following instructions for Diotic FM:

> "Now the sound modification will be slightly different. Instead of wobbling between your ears, one of the sounds will fade in and out. Can you detect the modified "special" sound?"

The following instructions for the Gap detection:

> "Next, each of the squares will carry two clicks so close to each other they sound like only one.  One of the squares in the middle will carry a pair of clicks with a bigger separation between them so they will sound slightly different. Your task is to identify the square with the different pair of clicks."

The second half of the Signal in Competition Battery contained the SRM assessments. Participants were shown the following instructions:

> "In each trial, you will hear a person call the name Charlie followed by the directions to go to a specific combination of color and number. Press the button on the grid that corresponds to such directions."

The remaining assessments contained similar instructions, and small breaks were provided between assessments. Each sub-battery would end with an instructions screen the following text or the equivalent:

> "You have completed this part of the evaluation, please let an experimenter know you are done :)".

An experimenter or research assistant would load the next sub-battery to continue testing or finish the session.
